# Divergent effects of a Treg-selective IL-2 mutein on Influenza specific T cell responses

**DOI:** 10.1101/2025.06.10.658962

**Authors:** Joseph R Albe, Anita Chaudhary, Asheema Khanna, Kristen N Weinstein, Steven F. Ziegler, Vandana Kalia, Surojit Sarkar, Daniel J Campbell

## Abstract

Enhancing regulatory T cell (Treg) function offers a compelling therapeutic strategy for autoimmune disease. Engineered IL-2 muteins selectively expand functional Tregs with minimal impact on other immune cells, but their potential to compromise antiviral immunity remains largely unexplored. Here, we used a murine model of Influenza A virus (Flu) infection to determine how IL-2 mutein shapes T cell responses to respiratory virus infection. IL-2 mutein administration prior to infection suppressed Flu-specific (Flu-sp) CD8 T cell responses and altered their localization and phenotype within the lungs, without affecting bystander CD8 T cells. This suppression correlated with reduced antigen presentation molecule expression on conventional dendritic cells (cDCs) early after infection but did not impact Flu-sp CD8 T cell priming. In contrast, administering IL-2 mutein during infection exacerbated disease and drove CD25-dependent expansion of Flu-sp CD8 T cells. Despite these opposing effects on effector responses, Fc.Mut24-treated mice generated robust antibody responses and protective T cell memory which were maintained for at least 170 days. These findings reveal that Fc.Mut24 has temporally distinct effects on antiviral immunity, dampening early effector responses when given before infection, but enhancing effector expansion and disease severity when delivered during infection. Our results provide critical context for the therapeutic application of IL-2 muteins and highlight the importance of treatment timing in balancing immune modulation with protective immunity.

## Introduction

Increasing regulatory T cell (Treg) abundance and function is a promising strategy for treating autoimmune diseases, as Tregs suppress inflammatory responses that drive disease pathology (1, 2). This can be accomplished by either expanding and re-infusing Tregs in various cell therapy approaches, or by boosting the abundance and/or function of endogenous Tregs. In the second category, one approach involves targeting the interleukin-2 receptor (IL-2R), which is critical for Treg development and function (3). The IL-2R exists in two distinct forms - a high-affinity form (composed of CD25, CD122, and CD132) is constitutively expressed by Tregs and activated CD4 and CD8 conventional T cells (Tconv), whereas the low-affinity IL-2R (composed of CD122 and CD132), is primarily found on resting naïve and memory Tconv and is highly expressed by NK cells. This differential expression of IL-2R forms helps support Treg homeostasis and function by allowing them to effectively compete for limiting amounts of IL-2 thereby preventing excessive immune activation (4). However, when IL-2 is produced in excess it can act on Tconv to support their differentiation into effector and memory populations (5). This receptor expression pattern also provides the rationale for low-dose IL-2 therapy, which aims to preferentially expand Tregs in autoimmune disease (6).

An emerging approach to more effectively target Treg cells for treatment of autoimmunity is to use engineered IL-2 <u>mut</u>ant prot<u>eins</u> (‘muteins’), which are IL-2 variants designed to enhance Treg selectivity (2, 7). Several Treg-biased IL-2 muteins have been developed, including the Fc-fused IL-2 mutein from Merck (MK-6194) and pegylated IL-2 from Nektar (Rezpegaldesleukin), each of which show enhanced Treg-selectivity and potent Treg expansion in pre-clinical models or early clinical trials (8, 9). Data from Rezpegaldesleukin trials have demonstrated efficacy in improving atopic dermatitis in Phase 1b (9), showing the potential of this therapeutic approach. We developed an Fc-fused murine IL-2 mutein (known as Fc.Mut24) with two amino acid substitutions that decreased CD122 binding and thus increased dependence on CD25 to induce functional signaling (10, 11). Although Fc.Mut24 is a weaker IL-2R agonist *in vitro* than an Fc-fused version of wild-type IL-2 (Fc.WT), it is highly Treg-selective and more effectively expands Tregs *in vivo*. In addition to increasing Treg abundance, Fc.Mut24 also enhances suppressive functions of Tregs, including CTLA-4-mediated transendocytosis of the co-stimulatory ligands CD80 and CD86 (12). Importantly, unlike Fc.WT, Fc.Mut24 can be administered at high doses while maintaining Treg selectivity and effectively prevented disease progression in the non-obese diabetic (NOD) mouse model of Type 1 Diabetes (10). Thus, Fc.Mut24 provides a novel tool for pre-clinical mechanistic studies of IL-2 mutein therapy.

Given their ability to potently expand Tregs and broadly suppress the immune system, IL-2 muteins represent a promising new class of autoimmune therapies. However, immunosuppressive treatments—including corticosteroids, IL-6 inhibitors, co-stimulation inhibitors, tumor necrosis factor (TNF)-α inhibitors—also impair immune responses to infections and reduce vaccine efficacy (13–17). While these therapies target specific inflammatory pathways, Tregs use multiple immunosuppressive mechanisms to dampen immune responses (18), and therefore therapies that augment Treg abundance and function may be even more detrimental for responses to infection. Respiratory virus infections, such as RSV, Influenza and SARS-CoV-2, are particularly relevant in this context, as several immunosuppressive therapies increase the risk of severe infection (19, 20), and Tregs play a role in dampening immune responses and promoting tissue repair during respiratory virus infections (21, 22), raising the possibility that IL-2 mutein therapy could increase the risk of poor outcomes. Influenza, in particular, remains a major global health burden causing significant morbidity and mortality each year. Given these risks, it is important to determine whether IL-2 muteins compromise antiviral immunity. In addition, IL-2-based therapies could also promote inflammation by activating T cells that upregulate CD25 and express the high-affinity IL-2R. Therefore, administering IL-2 mutein during infection, when T cells are already activated and have upregulated the high-affinity IL-2 receptor, could enhance signaling and drive excessive T cell activation, proliferation, and immunopathology. Thus how the various activities of IL-2 mutein combine to impact the outcome of respiratory virus infection is complex, and may depend on the timing of IL-2 mutein administration.

We used a murine model of Influenza A virus (Flu) infection to investigate how Fc.Mut24 influences antiviral T cell responses during respiratory infection. Administering Fc.Mut24 before infection suppressed Flu-specific (Flu-sp) CD8 T cells at 9 days post-infection (dpi) and altered their localization and phenotype within the lung. This suppression was specific to Flu-sp CD8 T cells, as bystander cells remained unaffected. Although Fc.Mut24 reduced antigen presentation molecule expression on conventional dendritic cells (cDCs) at 3dpi, the number of Flu-sp CD8 T cells was unchanged at 4dpi. In contrast, administering Fc.Mut24 during infection worsened disease severity and drove an expansion of Flu-sp CD8 T cells that was dependent on cell-intrinsic CD25 expression. Despite the divergent effects of Fc.Mut24 treatment before or during infection, Fc.Mut24-treated mice generated protective memory that persisted through at least 170dpi. These findings demonstrate that the immunologic context in which IL-2 mutein therapy is administered critically shapes antiviral T cell responses, highlighting the need to evaluate how immune-modulating treatments for autoimmunity may influence host defense.

## Results

### Fc.Mut24 treatment before infection suppresses Flu-specific CD8 T cell responses

In patients receiving IL-2 muteins for autoimmunity, administration of mutein and expansion of Tregs just prior to respiratory virus infection could dampen the immune response, worsen outcomes, and compromise development of durable memory against re-infection. To model this possibility, we gave 10 μg of Fc.Mut24 to female C57BL/6 (B6) mice one day before intranasal infection with 2000 plaque-forming units (PFU) of Influenza A virus (strain A/HK-x31(H3N2)) (X31). The relative timing of Fc.Mut24 treatment and infection aimed to maximize Treg expansion during the T cell priming phase to determine how this impacted the Flu-sp immune response. We assessed T cell responses in the lungs and spleen at 9dpi, corresponding to the peak of the effector T response in our model.

Consistent with its ability to potently expand Tregs, Fc.Mut24-treated mice exhibited a higher frequency and total number of Tregs in both lungs and spleen compared to vehicle controls (Figure 1A and Supplemental Fig. 1A). Flu-sp CD8⁺ T cells play a critical role in viral clearance, memory formation, and cross-protection against heterosubtypic influenza strains, and we therefore focused our analyses on the impact of Fc.Mut24 treatment on this population, using an MHC class I tetramer recognizing the immunodominant nucleoprotein antigen (NP)_366-374_ to identify Flu-sp cells. At 9 dpi Fc.Mut24-treated mice had a significant reduction in both the frequency and total number of Flu-sp CD8⁺ T cells in the lungs (Figure 1B) and spleen (Supplemental Figure 1B) compared to vehicle-treated controls. The lung is a complex organ that is highly vascularized, and T cells specific for respiratory pathogens can be found in both the lung parenchyma and patrolling the lung vasculature (23). Therefore, we used intravenous (IV) CD45 antibody labeling to distinguish Flu-sp T cells found in the lung vasculature (IV⁺) and lung parenchymal (IV⁻) Flu-sp CD8⁺ T cells. Fc.Mut24-treated mice had a significant enrichment of Flu-sp CD8⁺ T cells within the lung parenchyma compared to vehicle-treated controls (Figure 1C). However, despite this skewed distribution in the lung parenchyma, the total number of Flu-sp CD8⁺ T cells remained lower in both the lung vasculature (∼7-fold reduction) and parenchyma (∼3-fold reduction) compared to control animals (Figure 1C).

**Figure 1:**
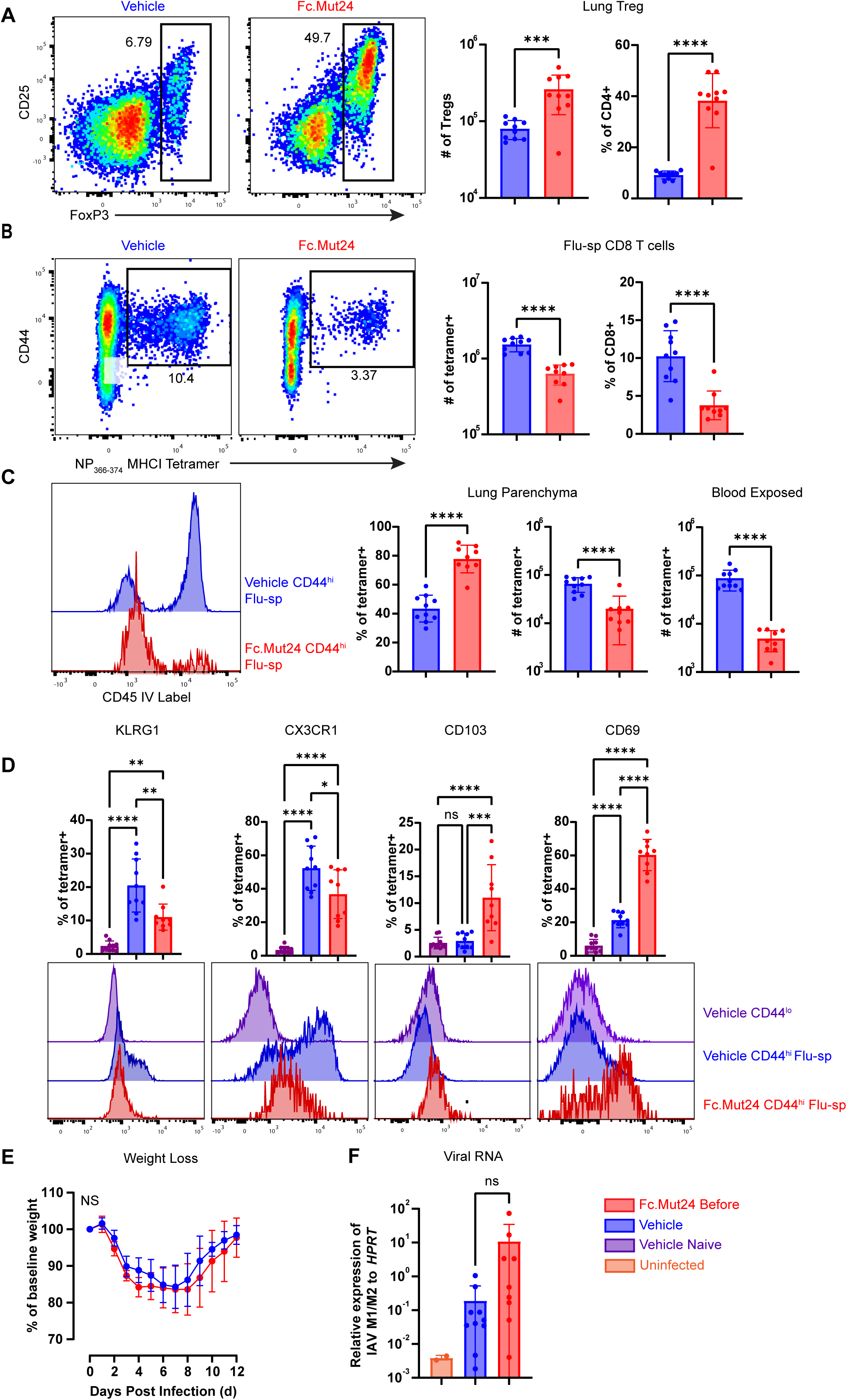
Fc.Mut24 administration before X31 infection limits Flu-sp CD8^+^ T cell responses. Mice were treated one day before infection with 2000PFU of X31 with either vehicle (blue) or Fc.Mut24 (red). All data represent takedown at 9dpi. **A)** Representative flow cytometry analysis of CD25 and Foxp3 expression by gated CD4^+^ T cells in the lungs of vehicle- and Fc.Mut24-treated mice (left), and quantification of the number and frequency of Tregs in the lungs (right). **B)** Representative flow cytometry analysis of CD44 expression and NP_366-374_ tetramer staining by gated CD8^+^ T cells in the lungs of vehicle- and Fc.Mut24-tretted mice (left), and quantification of the number and frequency of Flu-sp T cells in the lungs (right). **C)** Representative histogram of IV label staining on gated Flu-sp T cells in lungs of vehicle- and Fc.Mut24-treted mice (left), and quantification of the frequency and number Flu-sp CD8 T cells in parenchymal vs blood exposed compartments (right). **D)** Representative flow cytometry staining and quantitative analyses showing expression of the selected markers by gated CD44^lo^ naïve CD8^+^ T cells in the lungs of vehicle-treated mice, or by Flu-sp T cells in the lungs of vehicle- or Fc.Mut24-treated mice as indicated. **E)** Weight loss as a percent of baseline in vehicle- or Fc.Mut24-treated mice during the course of X31 infection. **F)** Quantification of viral genes *M1/M2* relative to *HPRT* by PCR in vehicle- or Fc.Mut24-treated mice as indicated. Each point represents analysis of an individual animal. All data are representative of two separate experiments *p < 0.05, **p < 0.01, ****p < 0.0001, by unpaired T test for A-C and one way ANOVA with multiple comparison for D and F and with mixed effects for E.

To further investigate the altered differentiation and migration of Flu-sp CD8⁺ T cells, we analyzed markers associated with tissue residency and vascular retention. Consistent with IV labeling results, the frequency of Flu-sp CD8⁺ T cells expressing KLRG1 and CX3CR1 (molecules linked to retention in the lung vasculature (24) was reduced in Fc.Mut24-treated mice, whereas expression of the tissue-residency markers CD103 and CD69 was increased (Figure 1D). This did not simply reflect their altered distribution, as Fc.Mut24-treated mice had higher expression of CD69 and CD103 compared with control mice even within the lung parenchyma fraction (Supplemental Figure 1C), and similarly showed significantly reduced expression of CX3CR1 and KLRG1 in the blood-exposed fraction (Supplemental Figure 1D) compared to the parenchymal compartment. To further determine how Fc.Mut24 treatment altered the functional differentiation of Flu-sp CD8^+^ T cells, we examined expression of transcription factors and cytokine receptors that promote effector and memory cell differentiation (Supplemental Figure 2A). Flu-sp CD8^+^ T cells from Fc.Mut24-treated mice showed a slight reduction in expression of the transcription factor EOMES, but no difference in expression of TCF1 or T-bet. However, consistent with increased memory potential, Fc.Mut24 treatment increased expression of the IL-7R component CD127 (25).

Despite the reduced magnitude and altered distribution and phenotype of the Flu-sp CD8⁺ T cell response, Fc.Mut24 treatment did not affect disease severity, as measured by weight loss (Figure 1E), viral RNA load (Figure 1F), or NP-specific IgG antibody titers (Supplemental Figure 1E) compared to vehicle controls. Notably, while Fc.Mut24 treatment did not alter the magnitude of the Flu-sp CD4⁺ conventional T cell response (Supplemental Figure 1F), there was a decrease in FoxP3^−^ CD4⁺ T cells in the lungs, suggesting a broader immunomodulatory effect (Supplemental Figure 1G). Further we observed no changes in the interferon gamma or IL-2 production by Flu-sp CD8 T cells that were stimulated *ex vivo* between vehicle treated and Fc.Mut24, suggesting there was no direct impact on functionality of the resulting response (Supplemental Figure 1H and I). Thus, Fc.Mut24 treatment resulted in sustained Treg expansion, and decreased Flu-sp CD8⁺ T cell abundance associated with altered localization and phenotype, with a larger fraction of cells localized to the tissue parenchyma, potentially positioning them for effective viral clearance. This is supported by the lack of change in disease severity or viral loads despite the overall reduction in Flu-sp CD8⁺ T cells.

### Fc.Mut24 Treatment Selectively Limits Flu-Specific Responses Without Suppressing Bystander CD8 T Cells

In addition to reducing the abundance and localization of NP-specific CD8^+^ T cells, Fc.Mut24 treatment also reduced the number of tetramer-negative CD44^hi^ (CD44^hi^) CD8^+^ T cells in the lungs at 9 dpi (Figure 2A), and these tetramer-negative cells also tended to localize more toward the lung parenchyma, although this trend did not reach statistical significance (Supplemental Figure 3A). These cells also showed a similar shift in expression of markers of T cell migration and functional differentiation as the NP-specific CD8^+^ T cell population (Supplemental Figure 3B). Ki-67 expression was also reduced in CD44^hi^ cells from Fc.Mut24-treated mice, suggesting decreased proliferation of these antigen-experienced cells (Supplemental Figure 3B). These tetramer-negative cells could represent Flu-sp cells recognizing epitopes other than the (NP)_366-374_ epitope, or alternatively could be bystander activated CD8^+^ T cells that also infiltrate the lungs during influenza infection (26, 27). To determine if Fc.Mut24 altered this bystander response, we infected mice intraperitoneally with 2000 PFU of Vesicular Stomatitis virus that expresses the SIINFEKL epitope from ovalbumin (VSV-OVA) and allowed 28 days for resolution. This generated SIINFEKL-specific CD8^+^ T cells, providing a bystander population that could be identified and tracked via staining with a K^b^/SIINFEKL MHCI tetramer. We then treated the mice with Fc.Mut24 or vehicle and, one day later, challenged half the mice intranasally with X31 or mock infection, creating four experimental groups for tracking the SIINFEKL-sp CD8^+^ T cell response. In contrast with the response of Flu-sp CD8^+^ T cells, we observed no significant differences in the number of SIINFEKL-sp CD8^+^ T cells in the lungs across all groups (Figure 2B). Although the frequency of SIINFEKL-sp was reduced in the VSV + Flu + vehicle group, this likely reflects the influx of Flu-sp CD8^+^ T cells (Figure 2B). Moreover, SIINFEKL-sp CD8^+^ T cells showed no differences in localization between the parenchyma and blood-exposed compartments across all groups (Supplemental Figure 3C). We also found no changes in the expression of selected markers that were altered on CD44^hi^ CD8^+^ T cells (TCF, CD127, CD69, Ki-67, KLRG1, or CX3CR1) on SIINFEKL - sp CD8^+^ T cells (Supplemental Figure 3D). Together, these results indicate that Fc.Mut24 does not suppress the SIINFEKL-sp response, suggesting that the observed suppression in the CD44^hi^ population likely reflects effects on Flu-sp cells that were not directly tracked, rather than non-specific bystander responses.

**Figure 2:**
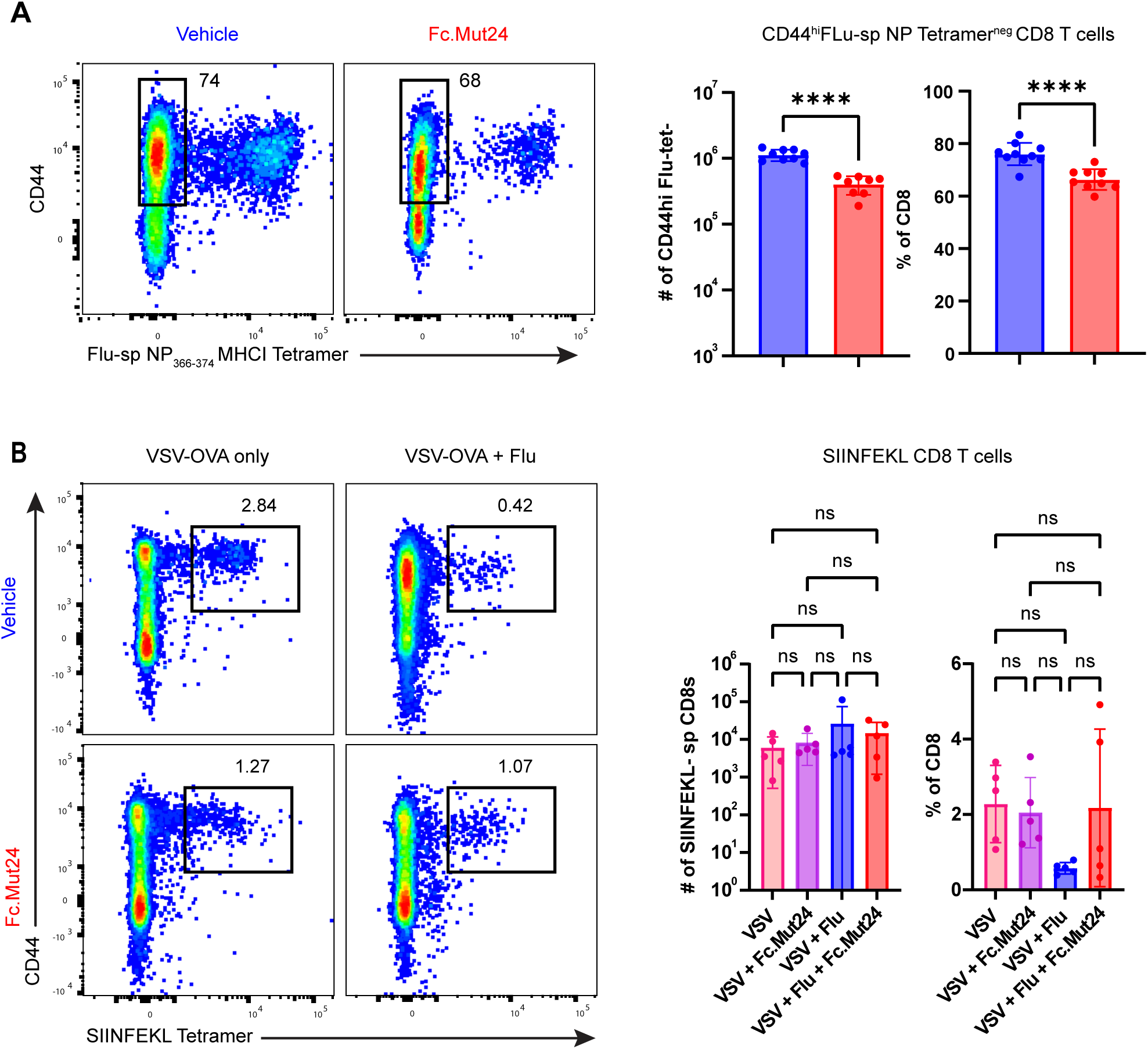
Fc.Mut24 treatment does not impact bystander CD8 T Cells. **A)** Mice were treated one day before infection with 2000PFU of X31 with either vehicle (blue) or Fc.Mut24 (red). Representative flow cytometry analysis of CD44 expression and NP_366-374_ tetramer staining by gated CD8^+^ T cells in the lungs of vehicle- and Fc.Mut24-tretted mice (left), and quantification of the number and frequency of CD44^hi^Tetramer^−^ T cells in the lungs (right). **B)** Mice were infected with VSV-OVA 30d before treatment with vehicle or Fc.Mut24 and X31 infection. All data represent takedown at 9dpi with X31. Representative flow cytometry analysis of CD44 expression and SIINFEKL tetramer staining by gated CD8+ T cells in the lungs of mice treated as indicated (left), and quantitative analysis of the number and frequency of SIINFEKL-sp CD8 T cells across groups (right). N = 5 per group. Each point represents analysis of an individual animal. *p < 0.05, **p < 0.01, ****p < 0.0001, by by unpaired T test for A and one way ANOVA with multiple comparison for B.

### Fc.Mut24 treatment prior to infection reduces cDC antigen presentation capacity with minimal impact on CD8 T cell proliferation at 4 days post-Influenza infection

Given the significant reduction in Flu-sp CD8^+^ T cells during the effector phase of the response, we wanted to determine if this was associated with changes in antigen presentation capacity of conventional dendritic cells (cDCs), which prime Flu-sp CD8^+^ T cell responses in the draining mediastinal lymph nodes (medLN) and spleen following Flu infection in mice (28). Indeed, by 3dpi, both cDC1 and cDC2 subsets strongly upregulated key antigen presentation markers, including CD80, CD86, and MHC I (Supplemental Figure 4A and B). We have previously shown that Tregs expanded through Fc.Mut24 upregulate CTLA-4 which targets co-stimulatory molecules CD80 and CD86 on cDCs, and that IL-10 production from expanded Tregs can modulate surface MHC expression (12). Therefore, to further assess how Fc.Mut24 affects Treg expansion and cDC activation in the context of Flu infection, we analyzed lung, medLN and spleen at 3 dpi. Fc.Mut24-treated mice showed a marked increase in FoxP3⁺ Tregs in both tissue sites (Figure 3A), which exhibited an effector phenotype defined by high expression of CTLA-4 and CD25 (Supplemental Figure 4C). Consistent with our previous results in LPS-treated mice and in NOD mice, the fraction of activated CD80^hi^CD86^hi^ cDC1 and cDC2 subsets in the lungs and spleen were significantly reduced in Fc.Mut24-treated mice at 3 dpi, whereas only cDC2 cells were significantly impacted in the medLN, (Figure 3B).

**Figure 3:**
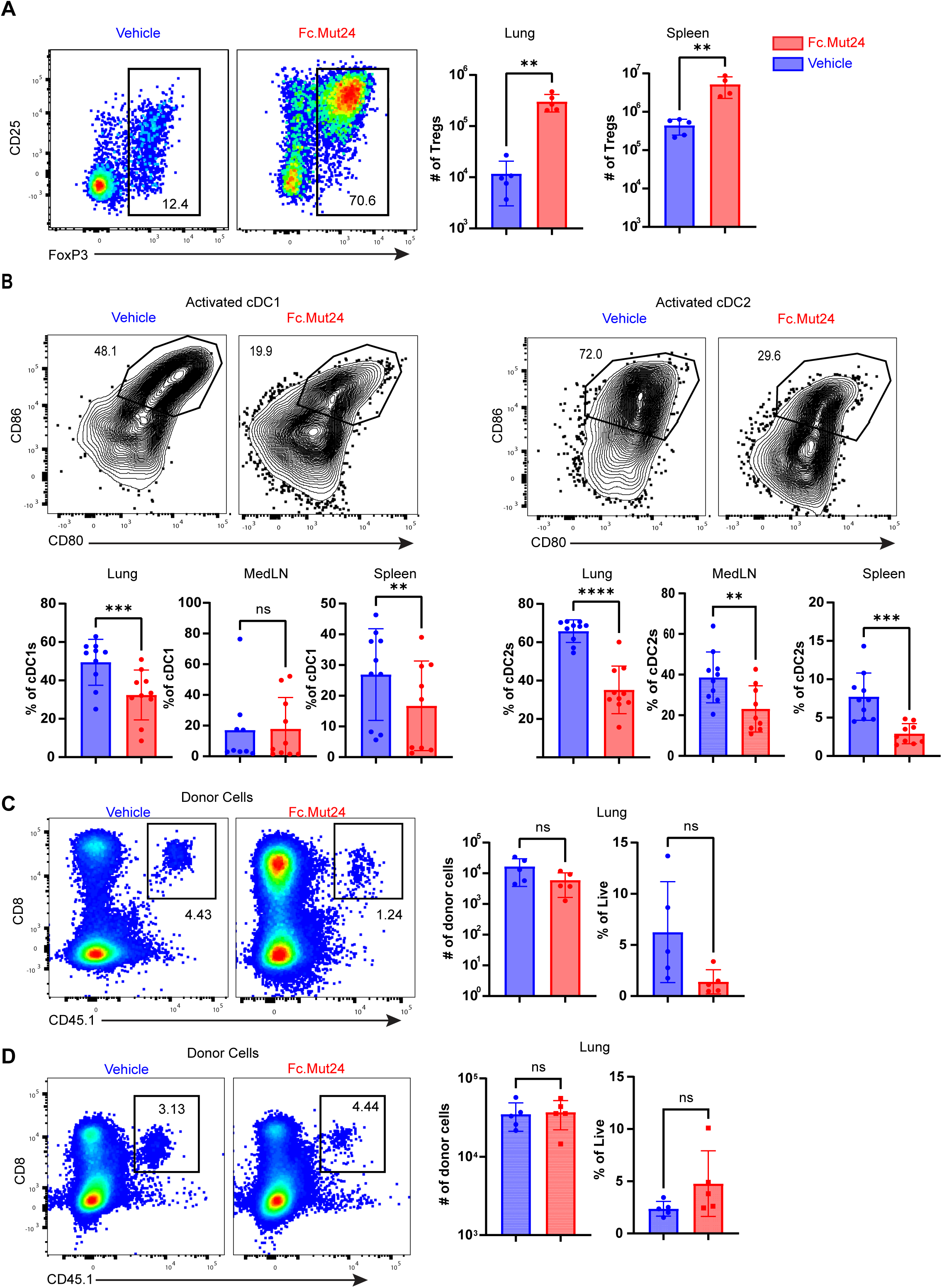
Fc.Mut24 treatment before X31 infection reduces antigen presentation capacity of DCs in the lungs and spleen. Mice were treated one day before infection with 2000PFU of X31 with either vehicle (blue) or Fc.Mut24 (red). **A)** Representative flow cytometry analysis of CD25 and Foxp3 expression by gated CD4^+^ T cells in the lungs of vehicle- and Fc.Mut24-treated mice (left), and quantification of the number and frequency of Tregs in the lungs (right) at 3dpi. **B)** Representative flow cytometry analysis of CD80 and CD86 expression by gated cDC1 (CD3/B220^−^Ly6G^−^F4/80^−^ MHCII^high^ CD11c^high^ XCR1^+^SIRPalpha^−^) and cDC2 (CD3/B220^−^Ly6G^−^F4/80^−^ MHCII^high^ CD11c^high^ XCR1^−^SIRPalpha^+^) cells from lungs at 3dpi with X31 (top). Quantification of activated CD80^+^CD86^+^ cDC1 or cDC2 in vehicle- or Fc.Mut24-treated mice as indicated. **C)** CD8^+^ T cells from CD45.1^+^CD45.2^+^ OT-I mice were enriched and transferred to CD45.2^+^ C57BL/6 recipients concurrent with vehicle- or Fc.Mut24-treatment one day prior to infection with X31-SIINFEKL. Representative flow cytometry analysis of CD8 and CD45.1 expression by gated live lymphocytes (left), and quantification of the frequency and number of transferred OT-I T cells (right) from the lungs of recipient mice at 4dpi (right). **D)** CD8^+^ T cells from CD45.1^+^CD45.2^+^ OT-I mice were activated for 48hrs and then transferred into transferred to CD45.2^+^ C57BL/6 recipients at 3dpi with X31-SIINFEKL. Representative flow cytometry analysis of CD8 and CD45.1 expression by gated live lymphocytes (left), and quantification of the frequency and number of transferred OT-I T cells (right) from the lungs of recipient mice (right) at 9dpi (6 days after cell transfer). Panels A, C and D are representative of one experiment n = 5 mice per group. Panel B is representative of two separate experiments n = 10 per group. Each point represents analysis of an individual animal. *p < 0.05, **p < 0.01, ****p < 0.0001, by unpaired T test for all panels.

Decreased antigen presentation capacity by cDCs could impair Flu-sp CD8⁺ T cell priming, resulting in the decreased expansion and infiltration of Flu-sp CD8^+^ T cells in the lungs we observed in Fc.Mut24-treated mice. To test this, we adoptively transferred congenically marked OT-I CD8⁺ T cells into Fc.Mut24- or vehicle-treated mice, then infected them with a recombinant X31 expressing the SIINFEKL peptide (X31-SIINFEKL). At 4 dpi, we measured OT-I expansion based on cell counts in the lung, medLN, and spleen. Surprisingly, although the number of OT-I cells trended lower in both the lungs (Figure 3C), spleen (Supplemental Figure 4D), Fc.Mut24-treated mice showed no significant reduction in OT-I expansion in any tissue, including the medLN compared to controls at 4dpi, and there were no significant differences in expression of the activation markers CD69, CD25, and CD44 in the lung or spleen (Supplemental Figure 4E). Thus, despite reducing co-stimulatory ligand expression by cDC2, pre-treatment with Fc.Mut24 had only minimal effects on the initial activation and tissue localization of virus-specific CD8^+^ T cells.

In addition to potentially impacting their initial priming, Fc.Mut24-mediated Treg expansion may suppress the migration and activation of Flu-sp CD8^+^ T cells within the lungs themselves. To test this, we activated OT-I CD8⁺ T cells *in vitro* with anti-CD3/CD28 Dynabeads for 48 hours to mimic T cell priming and then transferred them into recipient mice that had been pre-treated with vehicle or Fc.Mut24 and infected with X31-SIINFEKL three days earlier and sacrificed the mice for analysis 6 days later (9dpi). Although the frequency and number of transferred cells were reduced in the spleens of Fc.Mut24-treated mice, we observed no reduction in the number of OT-I cells in the lungs (Figure 3D, Supplemental Figure 4F). Additionally, rather than being suppressed the transferred OT-I cells showed increased expression of activation markers in the lungs but not the spleen (Supplemental Figure 4G). Together, these results indicate that while Fc.Mut24-driven Treg expansion dampens cDC activation and reduces the expansion and differentiation of Flu-sp cells, it has limited impact on their initial priming, and may even potentiate reactivation of virus-specific CD8⁺ T cells in the lungs.

### Fc.Mut24 administration during infection enhances Flu-specific CD8 T cell responses

A second important clinical scenario to model for how IL-2 mutein therapy may impact anti-viral immune responses is a patient receiving IL-2 mutein after initiation of infection, which would likely impact the Flu-sp T cell response differently than pre-infection treatment. These effects could include ‘off-target’ stimulation of CD25^+^ responding T cells leading to increased proliferation and effector cell differentiation that could enhance viral clearance but exacerbate detrimental immunopathology. However, Tregs also help coordinate tissue repair responses in the lungs during Flu infection, and this may help offset these inflammatory effects. Therefore, to model the effects of post-infection treatment, we administered Fc.Mut24 or vehicle to mice at 4dpi with X31 and analyzed immune responses at 9 dpi (five days after mutein treatment). Fc.Mut24 treatment at 4dpi robustly increased both the number and frequency of Tregs in the lung and spleen (Supplemental Figure 5A), confirming that it retained its ability to expand Treg even in highly inflammatory conditions as previously reported (10). However, rather than dampening the Flu-sp CD8^+^ T cell response, Fc.Mut24 given at 4dpi did not significantly change the frequency of Flu-sp CD8^+^ T cells, and increased their absolute number in the lung (Figure 4A) and spleen (Supplemental Figure 5B) at 9dpi. In contrast to results from mice pre-treated with Fc.Mut24, we observed a trend toward increased Flu-sp CD8 T cell frequency in the lung vasculature in mice given Fc.Mut24-after infection, and the absolute number of cells in this compartment was significantly increased whereas cell number in the lung parenchyma was not changed (Figure 4B). Fc.Mut24-treatment after infection did not alter expression of KLRG1, EOMES T-bet or TCF1, however similar to mice receiving Fc.Mut24 before infection, expression of CX3CR1 was reduced and CD127 expression was dramatically increased in Flu-sp CD8 T cells from the lungs of mice treated with Fc.Mut24 at 4dpi (Supplemental Figure 6A).

**Figure 4:**
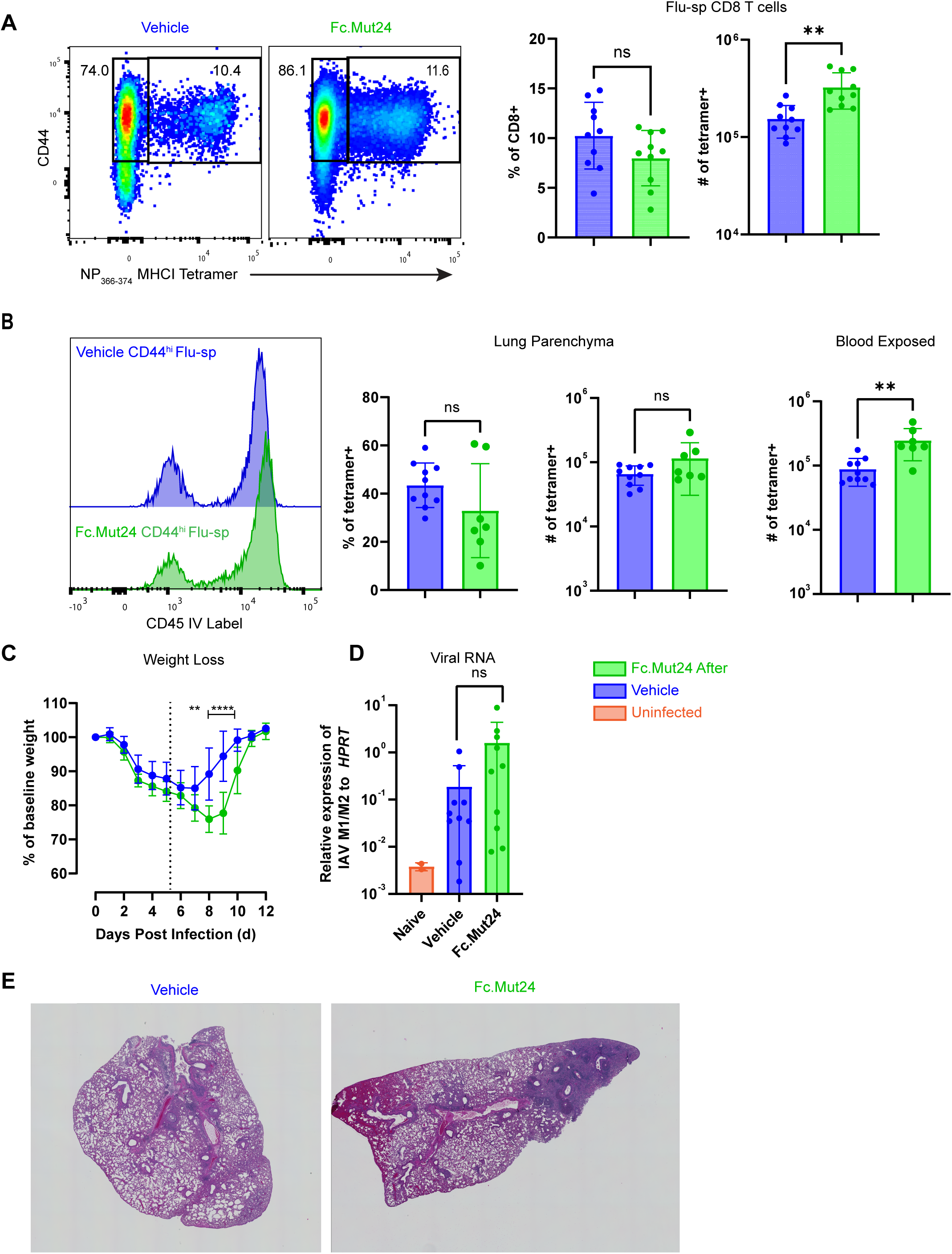
Fc.Mut24 treatment during infection enhances the Flu-specific CD8 T cell response. Mice were treated at 4dpi with X31 with either vehicle (blue) or Fc.Mut24 (green). All data represent takedown at 9dpi. **A)** Representative flow cytometry analysis of CD44 expression and NP_366-374_ tetramer staining by gated CD8^+^ T cells in the lungs of vehicle- and Fc.Mut24-tretted mice (left), and quantification of the number and frequency of Flu-sp T cells in the lungs (right) **C)** Representative histogram of IV label staining on gated Flu-sp T cells in lungs of vehicle- and Fc.Mut24-treted mice (left), and quantification of the frequency and number Flu-sp CD8 T cells in parenchymal vs blood exposed compartments (right). **C)** Weight loss as a percent of baseline in vehicle- or Fc.Mut24-treated mice during the course of X31 infection. Dotted line indicates time of vehicle or Fc.Mut24 treatment. **D)** Quantification of viral genes *M1/M2* relative to *HPRT* by PCR in vehicle- or Fc.Mut24-treated mice as indicated. **E)** Representative H&E images from mice at 12 days post infection. All data are representative of two separate experiments with n = 10. Each point represents analysis of an individual animal. *p < 0.05, **p < 0.01, ****p < 0.0001, unpaired T test for panels A and B and one way ANOVA with multiple comparison for D and Mixed Effects for panel C.

Unlike mice given Fc.Mut24 before Flu infection, mice treated at 4dpi lost more weight and recovered more slowly than vehicle-treated controls (Figure 4C). This was not associated with increased viral RNA in the lungs (Figure 4D) and thus is likely due to exacerbated immunopathology (Figure 4E). Importantly, this was not due to heightened functionality of Flu-specific CD8⁺ T cells, as they did not show increased IFN-γ or IL-2 production following ex vivo peptide stimulation (Supplemental Figure 5E and F). As we observed with Flu-sp CD8+ T cells, Flu-sp CD4+ T cell responses were also increased in mice given Fc.Mut24 at 4dpi (Supplemental Figure 5C). However, despite the ability of IL-2 to inhibit the differentiation of Tfh cells and subsequent antibody responses during infection (29, 30), NP-specific IgG titers were not altered by Fc.Mut24 treatment (Supplemental Figure 5D).

### Expansion of Flu-specific CD8 T cells by Fc.Mut24 requires CD25 expression

Fc.Mut24 requires CD25 to initiate IL-2R signaling, and activated STAT5 downstream of this pathway can further amplify CD25 expression through a positive feedback loop (31). We therefore hypothesized that in mice given Fc.Mut24 at 4dpi, CD25 expression on primed Flu-sp CD8^+^ T cells enables direct stimulation by Fc.Mut24, leading to enhanced expansion and altered differentiation and localization. Indeed, Flu-sp CD8^+^ T cells expressed higher levels of CD25 in 4dpi Fc.Mut24-treated mice, and thus Fc.Mut24 may directly stimulate responding CD8^+^ T cells and promote their excessive proliferation (Figure 5A). To directly test this hypothesis, we used CRISPR/CAS9 RNPs to delete the *Il2ra* gene (which encodes CD25) in naïve OT-I T cells (32), and successful targeting was confirmed by assessing CD25 upregulation in targeted cells after *in vitro* stimulation with anti-CD3/anti-CD28 (Supplemental Figure 7A). We transferred 5 × 10 cells to recipient CD45.2+ animals prior to infection with X31-SIINFEKL, and infected animals were treated with either vehicle or Fc.Mut24 at 4dpi. Indeed, loss of CD25 significantly impaired the Fc.Mut24-mediated hyper-expansion of OT-I cells following X31-SIINFEKL infection, whereas there was no significant difference in expansion of control and *Il2ra*-targeted cells in vehicle-treated mice (Figure 5B). We also analyzed the response of endogenous host-derived SIINFEKL-sp T cells using K^b^/SIINFEKL tetramers. Similar to what we observed for CD8^+^ T cells specific for the NP_366-374_ Flu epitope, Fc.Mut24 treatment led to an increased expansion of the endogenous SIINFEKL-sp cells compared to mice that received vehicle (Figure 5C), and thus the changes in expansion we observed are not epitope-specific, but rather reflect direct stimulation of CD25-expressing CD8 T cells by Fc.Mut24 during infection, which likely contributes to the increased presence Flu-sp CD8^+^ T cells and the elevated immune-pathology associated with Fc.Mut24 administration post-infection.

**Figure 5:**
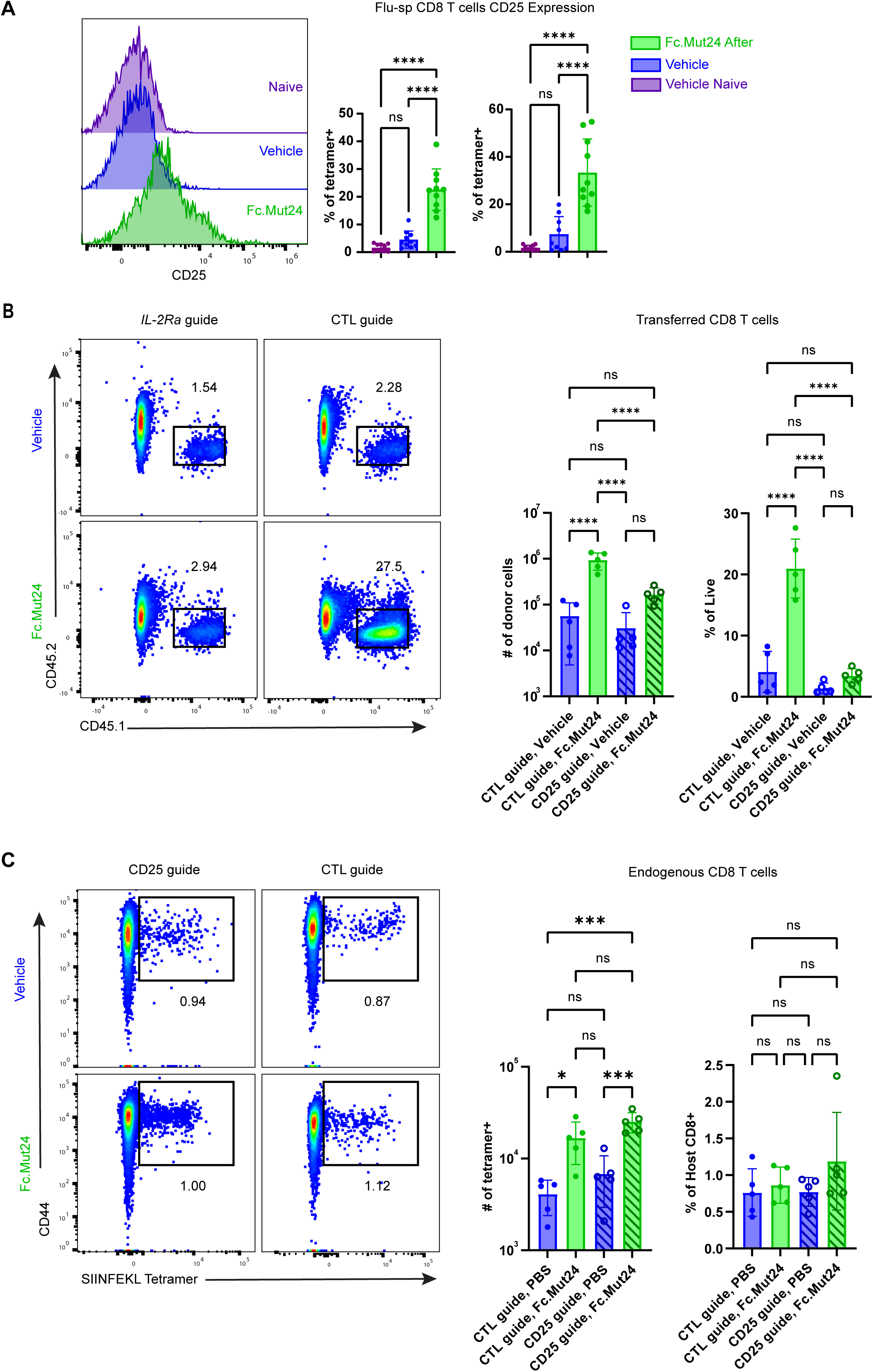
Fc.Mut24 directly expands IAV-specific CD8 T cells. 5×10^5^ congenic OT-I CD8 T cells were transferred to mice, that were a day later infected with X31-SIINFEKL and treated with 10ug of Fc.Mut24 at 4dpi. Data represents 9dpi takedown of lungs of indicated mice. **A)** Mice were treated at 4dpi with X31 with either vehicle (blue) or Fc.Mut24 (green). Data represent takedown at 9dpi. Representative flow cytometry staining and quantitative analyses showing CD25 expression by gated CD44^lo^ naïve CD8^+^ T cells in the lungs of vehicle-treated mice, or by Flu-sp T cells in the lungs of vehicle- or Fc.Mut24-treated mice as indicated. **B)** CD8^+^ T cells from CD45.1^+^ OT-I mice were enriched transfected with CAS9 RNPs containing a guide (g)RNA targeting the *Il2ra* locus or a control non-targeting gRNA. Cells were transferred to CD45.2^+^ C57BL/6 recipients that were infected with X31-SIINFEKL and treated with vehicle of Fc.Mut24 4 days later as indicated. Representative flow cytometry analysis of CD45.1 and CD45.2 expression by gated CD8+ T cells from the lungs of mice at 9dpi showing gate for donor OT-I T cells (left), and quantitative analysis of the abundance and frequency of donor OT-I cells (right). **C)** Representative flow cytometry analysis of CD44 expression and SIINFEKL tetramer binding by CD45.2+CD45.1-recipient cells (left) and quantitative analysis of endogenous SIINFEKL-sp CD8^+^ T cells from lungs of mice treated as in B. Data are from one experiment with n=5 mice/group, representative of two separate experiments. *p < 0.05, **p < 0.01, ****p < 0.0001, by one way ANOVA with multiple comparisons.

### Protective Memory formation and durability are maintained with Fc.Mut24 treatment

Mice treated with Fc.Mut24 before Flu infection showed reduced CD8^+^ T cell responses, which could impair memory formation and durability. Conversely, mice treated at 4dpi exhibited enhanced proliferation and effector phenotypes, which might also compromise memory development. However, regardless of treatment time, Flu-sp CD8^+^ T cells from Fc.Mut24-treated mice at 9dpi expressed elevated levels of the memory precursor marker CD127 which allows cells to respond to IL-7 and promotes memory cell formation. Therefore, we evaluated protective memory at 30 dpi. For this, we challenged previously infected mice with a lethal dose (5000 PFU) of the A/PR/8 (PR8, H1N1) (PR8) influenza strain. Importantly, PR8 (H1N1) and X31 (H3N2) are serologically distinct strains, and protection against this heterosubtypic infection is partially T cell dependent (33). All mice treated with Fc.Mut24 (either before or after initial X31 infection) were fully protected from PR8 infection at this timepoint, showing weight loss and recovery comparable to vehicle-treated X31 immune controls and surviving the challenge, whereas naïve mice all succumbed to PR8 (Figure 6A).

**Figure 6:**
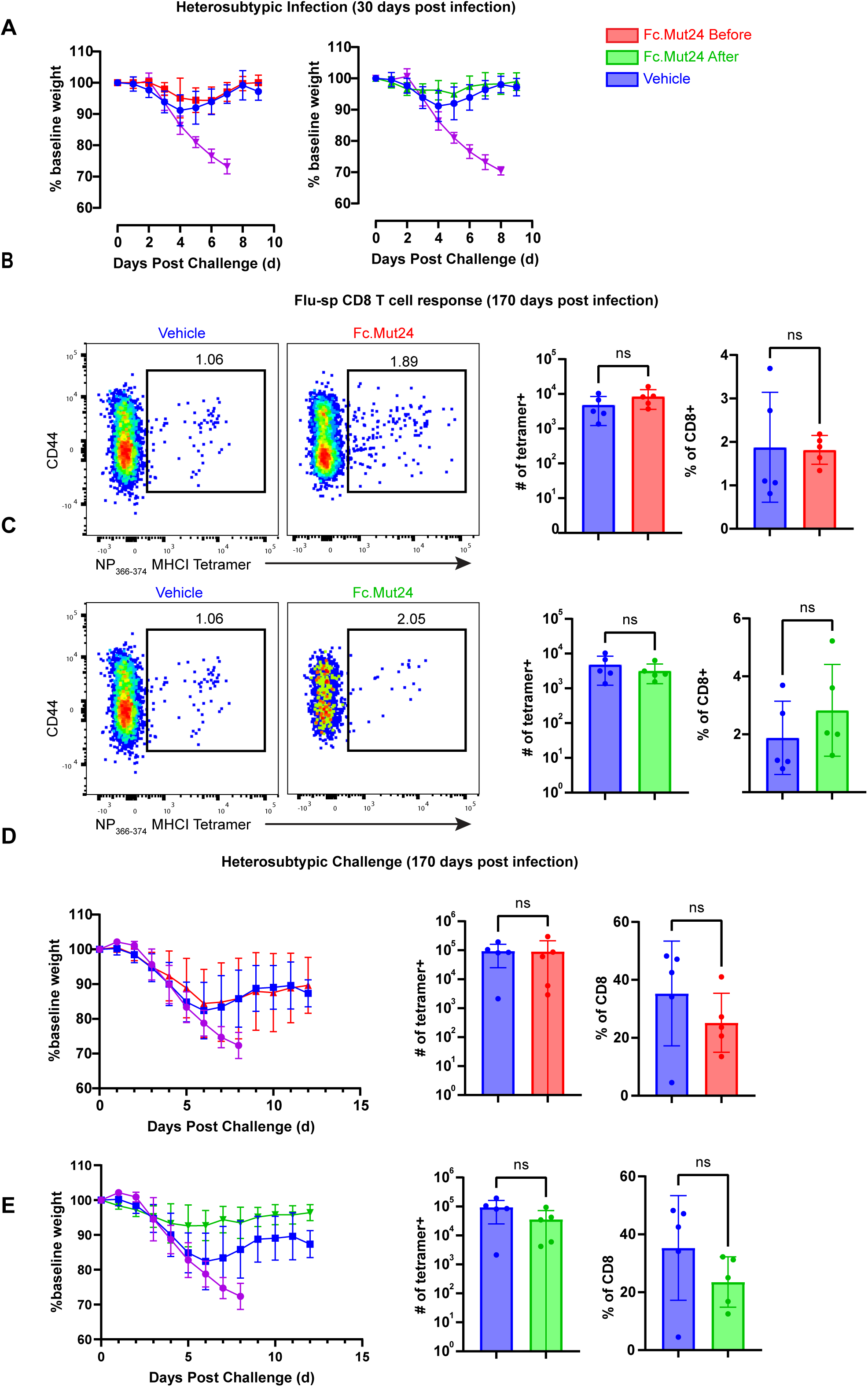
Fc.Mut24 does not alter generation or durability of protective T cell memory to influenza. Mice were infected with X31 and treated with vehicle (blue), or with Fc.Mut24 either a day before (red) or 4 days after (green) infection.. **A)** Mice were lethally challenged with 5000 FU of PR8 at 30 days post X31 infection. Disease severity based on weight loss from start of lethal challenge in mice treated with Fc.Mut24 one day before (left) or 4 days after (right) initial X31 infection (left) compared with vehicle-treated and X31-naïve mice. **B), C)** Representative flow cytometry analysis of CD44 expression and NP_366-374_ tetramer staining by gated CD8^+^ T cells in the lungs of vehicle- and Fc.Mut24-tretted mice, and quantification of the number and frequency of CD44^hi^Tetramer^−^ T cells in the lungs at 170dpi in mice treated one day prior to **B)** or 4 days after **C)** initial X31infection. **D), E)** X31-immune mice treated with vehicle or Fc.Mut24 were lethally challenged with 5000 FU of PR8 at 170dpi. Disease severity based on weight loss from start of lethal challenge (left). Quantification of Flu-sp CD8 T cells in lung (right) in mice treated with Fc.Mut24 one day prior to **D)** or 4 days after **E)** initial X31 infection. *p < 0.05, **p < 0.01, ****p < 0.0001, by unpaired T test and two way ANOVA with mixed effects for weight loss curves in panels A, C, D and E.

To assess long-term durability of protective memory, we treated mice with Fc.Mut24 either one day before or 4 days after X31 infection, and evaluated memory T cell abundance and protective responses at 170 dpi. Flu-sp CD8^+^ T cells remained detectable in the lungs of all animals, and both their frequency and number were comparable between Fc.Mut24- and vehicle-treated groups (Figure 6B and C). We also examined localization of memory cells in the lung parenchyma vs. vasculature, and expression of compartment-specific residency markers (CD69, CX3CR1, KLRG1), but found no significant differences at 170 dpi in any of our experimental groups (Supplemental Figure 8A and B). Upon lethal PR8 challenge infection, both vehicle- and Fc.Mut24-treated X31 immune were protected from lethal disease, and mice treated with Fc.Mut24 at 4dpi during initial X31 infection trended toward milder disease and less weight loss compared to vehicle controls, although this difference did not reach statistical significance (Figure 6 D and E). Moreover, the Flu-sp CD8^+^ T cell response showed no differences in magnitude, phenotype, or tissue localization between any of the X31 immune groups (Figure 6E; Supplemental Figure 8C and D). Thus, although Fc.Mut24 alters the acute CD8 T cell response—either suppressing or enhancing expansion of Flu-sp CD8^+^ T cells depending on timing—it does not compromise the generation or maintenance of protective T cell memory. Similarly, there were no differences in the maintenance of IgG responses to NP at 170dpi (Supplemental Figure 8E), and thus our findings show that regardless of timing relative to initial infection, Fc.Mut24 treatment does not impact the integrity and durability of memory Flu-sp CD8^+^ T cell responses.

## Discussion

Our study demonstrates that the timing of IL-2-based immunotherapy critically shapes its impact on anti-viral CD8⁺ T cell responses. Using Fc.Mut24, a Treg-selective IL-2 mutein, we show that treatment before respiratory virus infection suppressed the expansion and function of Flu-specific CD8⁺ T cells and alters their localization within the lungs, whereas treatment after infection expands virus-specific CD8^+^ T cells through CD25-dependent signaling and worsens symptomatic disease, likely through enhanced immunopathology. However, regardless of timing of treatment, mice recovered from infection and developed durable protective immunity against lethal heterosubtypic infections. These divergent and context-dependent effects appear to reflect a balance between suppressive Treg activity, and direct IL-2 signaling that promotes differentiation of CD25⁺ effector CD8⁺ T cells.

Administering Fc.Mut24 before infection significantly reduced the abundance of Flu-specific CD8⁺ T cells. Interestingly, despite this overall reduction, these cells showed a skewed localization toward the lung parenchyma. Viral load and disease severity remained unchanged despite the reduced number and altered distribution of CD8^+^ T cells, and this indicates that even in the face of massive Treg expansion, it is possible to generate protective Flu-specific CD8⁺ T cell responses to efficiently clear the infection. Through expansion of Treg cells expressing high levels of CTLA-4 Fc.Mut24 treatment before infection likely limits the availability of the CD80 and CD86 co-stimulatory ligands. Similarly, the large reservoir of CD25^hi^ Treg cells may reduce IL-2 availability for Flu-sp CD8^+^ T cells during the expansion phase, and together these factors may favor the priming and expansion of CD8⁺ T cells that receive strong TCR signals and effectively clear virus and form protective memory despite the immunosuppressive environment present during priming. Indeed, their function as an IL-2 sink is a key mechanism by which Treg can suppress CD8^+^ T cell responses (34, 35), and this takes place in a key priming window that occurs 2-3 days after initial activation (36). Since IL-2 signaling directly inhibits CD127 expression by responding CD8^+^ T cells (37), this could also help explain why formation of durable heterosubtypic immunity was intact despite a reduced initial effector response. IL-2 signaling during priming also drives CX3CR1 expression (38), and decreased IL-2 availability in Fc.Mut24 pre-treated mice may therefore decrease expression of this receptor and promote parenchymal rather than vascular T cell localization in the lung. Despite limiting expansion of Flu-sp CD8⁺ T cells, Fc.Mut24 had minimal effects on bystander T cells during X31 infection. As the recruitment and activation of bystander CD8^+^ T cells is driven largely by chemokines and cytokines produced by cells of the innate immune system (39), this supports a mechanism whereby Treg cells interrupt key pathways central to elaboration of the antigen-specific adaptive immune response.

Interestingly, we also observed no differences in total IgG specific for the viral NP protein at any time point examined. This is surprising given that IL-2 is a key inhibitor of Tfh differentiation and function (29, 30), and therefore restricting IL-2 availability by expanding Treg prior to infection may be expected to promote better antibody responses. Indeed, despite the reduced CD8⁺ T cell response we observed in Fc.Mut24 pre-treated mice, these animals successfully cleared the virus and established durable protective memory that was quantitatively indistinguishable from that observed in vehicle-treated mice, indicating that long-term memory formation and maintenance was not strictly dependent on the magnitude of the initial clonal expansion.

When administered 4 days after infection had begun, Fc.Mut24 drove hyper-expansion of Flu-sp CD8^+^ T cells. Under these conditions, we found that CD25-dependent signaling in responding Flu-sp CD8^+^ T cells promoted their increased expansion and accumulation in the lungs. We hypothesize that this results in enhanced immune-mediated pathology in the lungs (40), consistent with the increased weight loss observed in these mice. Interestingly, as we observed with Fc.Mut24 pre-treatment, flu-specific CD8^+^ T cells in mice given Fc.Mut24 four days after infection also expressed elevated CD127 and decreased CX3CR1, although this did not result in enhanced parenchymal localization. The direct impact of IL-2 on CD8^+^ T cell differentiation is thought to take place in a distinct priming window that would occur before Fc.Mut24 treatment in our system (36), and therefore we hypothesize that the changes in receptor expression we observe are due to enhanced proliferation of CD127^+^CX3CR1^−^ memory precursor cells in response to Fc.Mut24 after this differentiation window has closed. In this way, Fc.Mut24 treatment during infection may actually boost lasting protective memory formation, as mice treated as such trended towards less weight loss than vehicle controls upon heterosubtypic PR/8 rechallenge. Consistent with this, others have also described IL-2 administration as boosting CD8^+^ memory T cell generation and function in the context of active inflammation (41, 42). Sustained antibody responses may be explained by the enhanced proliferation of Tregs which have been shown to favor Tfh generation during influenza infection (43), thus enabling sufficient antibody responses and explain why we did not see reduced total IgG against the viral NP.

Together, our findings demonstrate that although treatment with Fc.Mut24 exerts immunosuppressive effects through Treg expansion, respiratory viral infection is still effectively controlled and durable memory responses are established, even when peak Treg expansion coincides with the initial priming phase. This observation has important implications for use of IL-2 muteins and other Treg-expanding immunotherapies, suggesting that the presence of Treg-mediated suppression does not necessarily preclude effective anti-viral T cell responses, viral clearance and development of protective immunity. However, these results also highlight a critical caveat in that IL-2-based therapies may exacerbate immunopathology if administered during active infection. This is especially relevant in the context of respiratory viral infections, where immune-driven tissue damage significantly contributes to morbidity and mortality (44). These results underscore the need for careful consideration of the timing and context of Treg-targeted interventions in infectious disease settings, particularly when balancing immune regulation with the necessity of effective pathogen clearance.

## Materials and Methods

### Mice

C57BL/6 (B6) (JAX Strain# 000664) mice were purchased from The Jackson Laboratory. B6.SJL-Ptprca Pepcb/BoyJ (CD45.1^+^, JAX Strain# 002014) and C57BL/6-Tg(TcraTcrb)1100Mjb/J (OT-I, JAX Strain # 003831) mice were bred and maintained at Benaroya Research Institute (BRI). Female Mice used in experiments between the ages of 6-10 weeks at start of experiment. Female mice are used due to male B6 mice being more resistant to infection (45). For histological analyses, euthanasia was performed with tribromoethanol overdose. For all other experiments mice were euthanized by CO_2_ inhalation. All experiments were performed in accordance with the guidelines and with the approval of the Institutional Animal Care and Use Committee of the BRI.

### Viruses and infections

Influenza viruses A/HK-x31(x31, H3N2), A/PR/8 (PR8, H1N1) (46), and the X31-ova expressing the CD8 H2-K^b^ restricted SIINFEKL epitope (47), were all generously provided by Dr. Kimberly D. Klonowski (University of Georgia). Animals were infected with 2000 PFU of either X31 strain or 5000PFU of PR8 strain intranasally in 50uL of PBS. All mice infected with Influenza were monitored at least daily for weight loss and twice a day when mice reached 80% of baseline weight. If mice reached 70% of baseline weight, mice were promptly euthanized using aforementioned methods. For longer term studies, once mice recovered they were monitored twice weekly. VSV-OVA (48) was generously provided from Dr. Pamela Fink (University of Washington). Mice were infected with 2000PFU of VSV-OVA, intraperitoneally in 100 uL of PBS.

### IL-2 mutein treatment

The development of the murine Fc.IL-2 mutein (Fc.Mut24) was previously described (10). Fc.Mut24 was produced and purified by Olympic Protein Technologies (Seattle, WA), and contained less than 15 endotoxin units (EU)/mL. For all experiments, 10 μg of Fc.Mut24 or PBS vehicle was administered via intraperitoneal injection.

### Cell isolation and flow cytometry

Where applicable, mice were anesthetized using 4% isoflurane and intravenously injected with 3μg of anti-CD45 or anti-CD45.2 antibody in 100uL for intravascular labeling. Mice were allowed to recover for two minutes before euthanasia, performed as described above. For flow cytometry, lungs were minced and digested in RPMI supplemented with Liberase (50 μg/mL) and DNase I (10 U/mL) for 20 minutes at 37 °C with agitation. Cell suspensions were filtered through 70 μm strainers into RPMI containing 10% FBS (RPMI-10), followed by red blood cell lysis with ACK lysis buffer and washing in RPMI-10. The same protocol was used for lymphoid tissues (mediastinal lymph node and spleen) when analyzing conventional dendritic cells (cDCs). Otherwise, spleens and medLNs were mechanically dissociated through a 70 μm strainer, followed by red blood cell lysis and washing as above. For dendritic cell analysis, CD11c⁺ cells were enriched using anti-CD11c microbeads (Miltenyi Biotec) according to the manufacturer’s instructions. Enriched cells were stained with selected antibodies targeting cDC surface markers (see reagent table).

For detection of antigen-specific CD4⁺ T cells, MHC class II tetramer staining was performed using the I-A^b^/NP_311-325_ (QVYSLIRPNENPAHK) tetramer (49), kindly provided by Marion Pepper (University of Washington). Samples were incubated with tetramer at room temperature for 1 hour, washed, and enriched using anti-APC microbeads (Miltenyi Biotec). Enriched cells were then stained with the CD4 flow cytometry panel, while the flow-through was used for additional staining and analysis. For CD8⁺ T cell analysis, H-2D^b^/NP_366-374_ (ASNENMETM) or H-2K^b^/SIINFEKL tetramers (generated by the NIH Tetramer Facility, Emory University) were used. Cells were stained with tetramer for 1 hour at room temperature prior to surface staining.

Cell surface staining for flow cytometry was performed in FACS buffer (PBS-2% BCS) using a selected antibody cocktail (See reagent table). Cells were incubated in the antibody cocktail for 20 min at room temperature and then washed in FACS buffer before collecting events on an BD Symphony flow cytometer. For intracellular staining, surface antigens were stained before fixation and permeabilization with FixPerm buffer (eBioscience) (See reagent table). Cells were washed and stained with selected antibodies. Flow cytometry data was analyzed using FlowJo software.

### Cell stimulations

Single-cell suspensions were prepared from tissues (as described above) and cultured in complete RPMI in the presence of influenza NP_366–374_ peptide (ASNENMETM, H-2D restricted) at 10 microMolar in 200 µL of cRPMI. Cells were incubated with secretion inhibitors (brefeldin A and monensin at 1x manufactures recommendation) throughout the stimulation period. Following stimulation, cells were stained with a viability dye and fluorochrome-conjugated antibodies against surface markers, then fixed, permeabilized, with BD cytoperm/Fix (cat. 554722) and stained for intracellular cytokines IL-2 and IFN-γ.

### Quantification of viral RNA

Total RNA was extracted from lung tissue using the Qiagen RNeasy Plus Mini Kit (cat. 74134) following the manufacturer’s protocol. Tissue homogenization was performed in RLT buffer containing β-mercaptoethanol (BME), with buffer volume adjusted according to tissue weight as specified by the kit. Approximately 15 mg of tissue was used per sample to avoid overloading the RNA-binding column. Samples were lysed using 5 mm stainless steel beads (Qiagen, cat. 69989) in 2.0 mL Safe-lock Eppendorf tubes on a TissueLyser II (Qiagen) at 30 Hz for 2 minutes. RNA yield was quantified using a NanoDrop spectrophotometer. Complementary DNA (cDNA) was synthesized from purified RNA using SuperScript II Reverse Transcriptase (Invitrogen, cat. 18064-014) and IDT ReadyMade Random Hexamer primers (cat. 51-01-18-26), including a no-RNA negative control in each batch. Reactions were assembled using a two-step thermal protocol. First, Master Mix 1 was added and samples were incubated at 65 °C for 5 minutes, then cooled to 4 °C. After addition of Master Mix 2, samples were incubated at 30 °C for 10 minutes followed by 42 °C for 50 minutes. The resulting cDNA was quantified using a NanoDrop spectrophotometer. Quantitative PCR (qPCR) was used to quantify Influenza A virus matrix (M1/M2) gene expression from PR8 and X31 strains. Reactions were prepared using IDT PrimeTime Std qPCR Assay containing a fluorescent probe and primers specific to the M segment of IAV with the following primer and probe design: Probe: (6-FAM/ZEN/IBFQ) 5′-CCTCTGCTGCTTGCTCACTCGATC-3′; Forward Primer: 5′-CAGCACTACAGCTAAGGCTATG-3′; Reverse Primer: 5′-CTCATCGCTTGCACCATTTG-3′. The probe/primer mix was resuspended at a 20x concentration. Each qPCR reaction (20 μL total volume) contained: 10 μL 2x IDT PrimeTime Gene Expression Master Mix (cat. 1055770),1 μL 20x PrimeTime qPCR Assay Mix, 4 μL diluted cDNA (∼75 ng), 5 μL nuclease-free water. An internal control targeting hypoxanthine-guanine phosphoribosyltransferase (HPRT) was run in parallel to normalize viral gene expression. The Viia7 Real-Time PCR System (Thermo Fisher Scientific) was used for amplification and detection.

### Adoptive transfers

Spleen and lymph nodes were harvested from OT-I mice, passed through a 70 μm strainer and ACK lysed. CD8^+^ cells were enriched by negative selection using a CD8 T cell isolation kit according to the manufacturer’s instructions (Miltenyi). Cells were then enumerated, washed in PBS before transfer by retroorbital injection into recipient mice (0.005-0.5×10^6^ cells per mouse depending on the experiment). For transfer of activated OT-I CD8 T cells, spleens were collected from CD45.1/.2 mice and processed as described above. Once in single cell suspension, 1×10^6^ cells were plated in a 12-well plate (see reagent table) in 1mL of cRPMI supplemented with 50UI/mL of rIL-2 (see reagent table). Cells were plated at a ratio 1:1 ratio with washed Dynabeads and allowed to culture for 48hrs. Following 48hrs, Dynabeads (see reagent table) were extracted using a magnet and cells washed before being transferred retroorbitally to mice at 0.5×10^6^ cells per mouse.)

### CRISPR deletion of Il2ra

For deletion of the *Il2ra* gene in primary naive OT-I TCR T cells, we utilized the CRISPR/CAS9 system as previously described (32). Purified CAS9 protein, *Il2ra* crRNA (AGAUGAAGUGUGGGAAAACGGUUUUAGAGCUAUGCU), and trans-activator RNA (tracrRNA) were purchased from Integrated DNA Technologies. CD8^+^ T cells were isolated from OT-I mouse spleen using CD8^+^ T cell negative selection MojoSort Kit (Biolegend). T cells were cultured in IL-7 (10ng/ml) for 24 hrs prior to transfection. CRISPR/CAS9 reagents were prepared as per the manufacturer’s instructions for IDT Alt-R CRISPR-CAS9 system. The pre-cultured CD8^+^ cells were were resuspended in T buffer (Invitrogen MPK1096) and electroporated using 6 2200V, 10ms, 3 pulses, with the Neon electroporation system. Following transfection, cells were transferred into warm RPMI with 10% FCS and were incubated at 37°C, 5% CO2 for 4-5 hours. Prior to adoptive transfer, the electroporation efficiency was evaluated by flowcytometry measuring ATTO uptake. Control cells were transfected with ATTO and CAS9 alone without *Il2ra* crRNA. To validate *Il2ra* deletion efficiency, the neon-transfected cells were plated on αCD3/αCD28 coated plates for activation for 48 hours followed by flow cytometry analysis of cell surface CD25 expression.

### Enzyme-Linked Immunosorbent Assay (ELISA)

To quantify serum antibody responses against influenza PR8 nuclear protein, ELISA plates were coated overnight at 4°C with anti-His antibody (1:1000 dilution in PBS, 50 μL/well). Plates were then washed three times with 100 μL PBS containing 0.1% Tween-20 (PBS-T), followed by incubation with His-tagged PR8 nuclear protein (SinoBiological, Cat# 11675-V08B) at 1:100 dilution in PBS-T (50 μL/well) for 1 hour at 37°C. After three additional PBS-T washes, wells were blocked with 100 μL of blocking buffer (PBS-T supplemented with 5% BSA) for 1 hour at 37°C. Diluted serum samples were added to the plates (50 μL/well) and incubated for 1 hour at 37°C. Plates were then washed three times with PBS-T and incubated with 50 μL of biotin-conjugated anti-mouse IgG antibody (Jackson ImmunoResearch, Cat# 115-035-062) diluted 1:1000 in blocking buffer for 1 hour at 37°C. After five washes with PBS-T, 50 μL of 1× TMB substrate was added to each well and color development was monitored. The reaction was stopped with 50 μL of H₂SO₄, and absorbance was measured at 450 nm using a microplate reader.

### Histological analysis

Lungs were excised and immediately fixed in 10% formalin for at least 24hrs and then paraffin embedded. H&E staining was performed on a 5μm tissue sections by Benaroya Research Institute Histology Core. Resulting slides were imaged using a Molecular Devices ImageXpress Confocal with the 10x objective by Benaroya Research Institute Histology Core.

### Statistics

All data are presented as mean values ± SD, and graphs were created and analyzed using Prism Software (GraphPad). Comparisons between treatment groups were analyzed using two-tailed unpaired student T tests, one-, or two-way ANOVA where appropriate, adjusted for multiple comparisons using Tukey’s post-test. To compare weight loss between experimental groups over time, a two-way mixed-effects ANOVA was performed. This mixed-effects approach allows for modeling the correlation structure of repeated observations and provides more accurate estimates by adjusting for inter-subject variability.

### Writing disclosure

Portions of the manuscript text were edited for grammar and clarity using OpenAI’s ChatGPT (version GPT-4). The tool was used solely for language refinement; all scientific content and interpretations are the original work of the authors.

## Supporting information

Reagent Table

## Acknowledgements

We would like to thank Pam Johnson and Riley Snodgrass for assistance with histology; Adam Wojno and Adin Pierce for help with flow cytometry; Andrew Biurch and the BRI veterinary staff for support with mouse work; and Marion Pepper, Mikel Ruterbusch, Kurt Pruner, and Geoff Hutchinson for providing the CD4 T cell tetramer. This work was supported by grants R01AI154773 and R21AI172140 from the NIH to D.J.C.

## Notes

### Competing Interest Statement

The authors have declared no competing interest.

### Summary of Updates

We have included new data assessing cytokine production by anti-viral CD8+ T cells in flu-infected mice treated with Fc.Mut24 (Figs S1H,I, and S5E, F), have expanded our discussion of key findings, and edited the some of the figures for clarity

