## Supplemental Figures 1-8 for "Divergent effects of a Treg-selective IL-2 mutein on Influenza specific T cell responses"

### Supplemental Data Figure 1

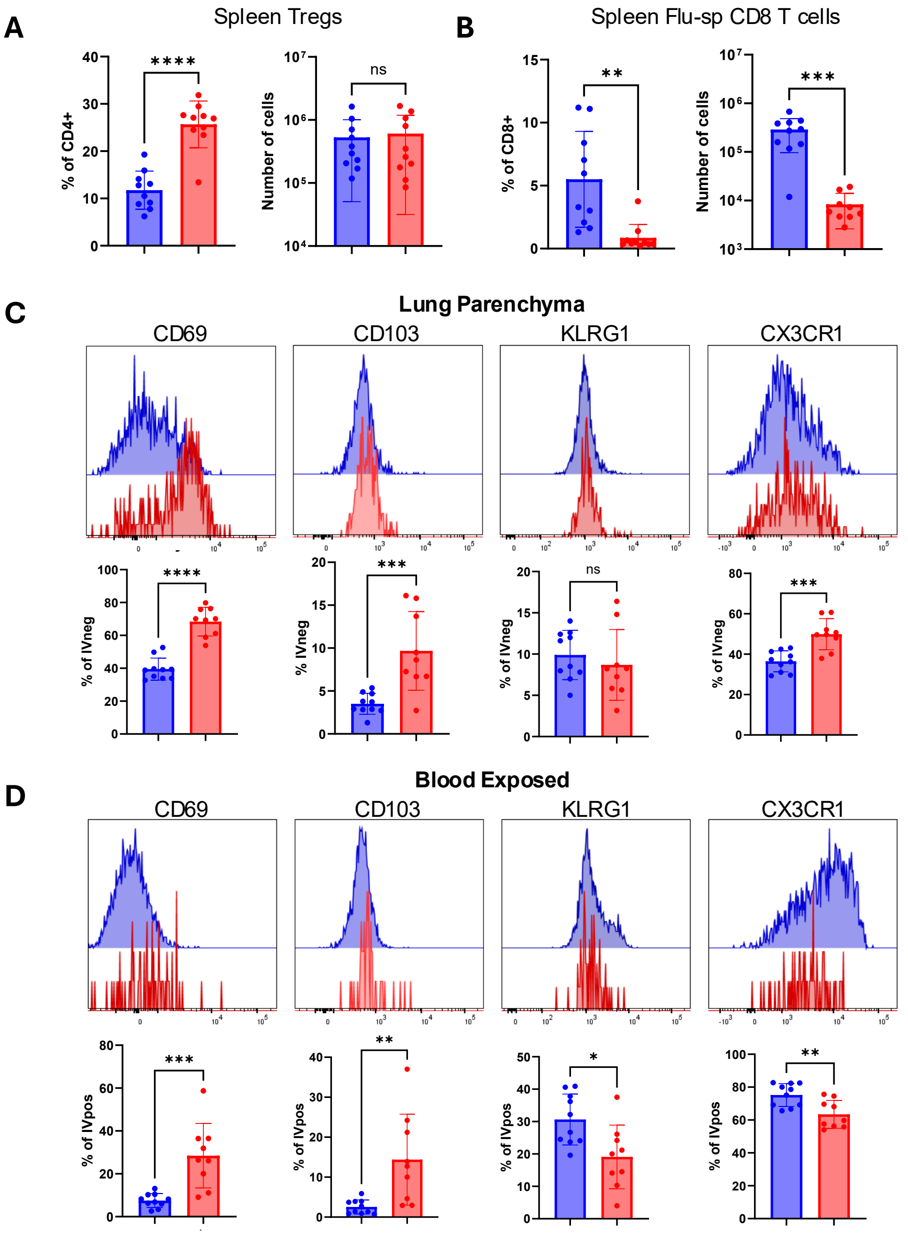


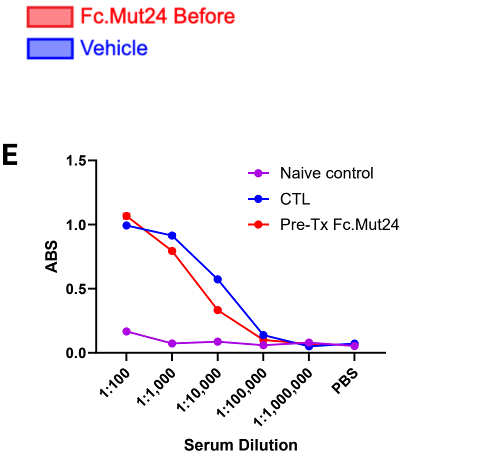


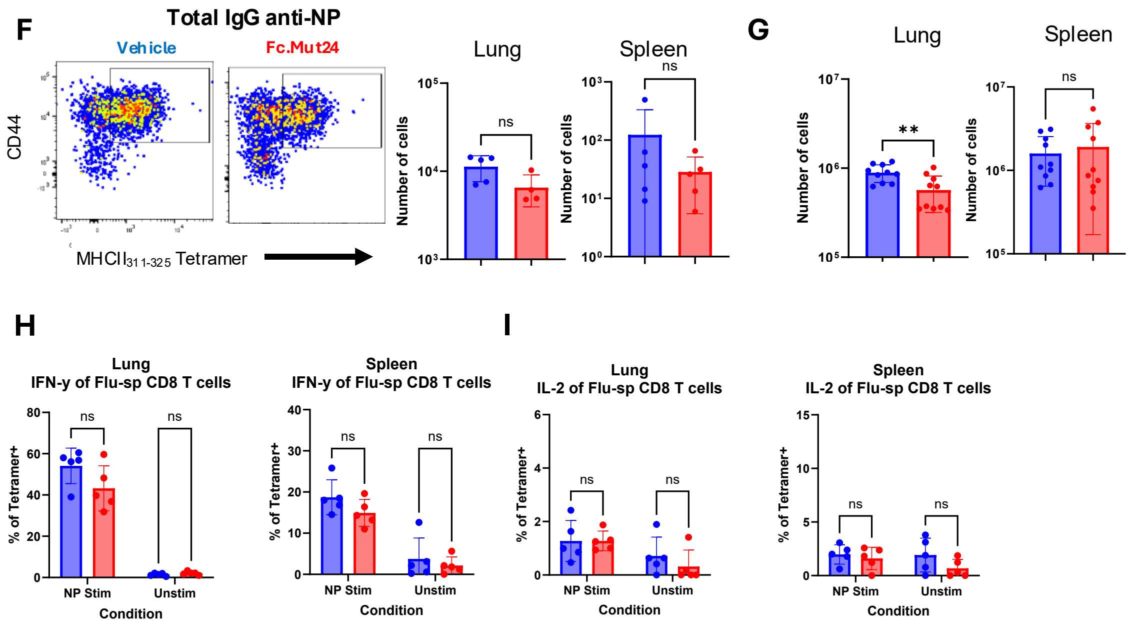


**Supplemental Figure 1.** Mice were treated with vehicle (blue) or Fc.Mut24 (red) one day before infection with 2000PFU of X31. All data represent takedown at 9dpi. **A), B)** Quantification of CD4$^+$Foxp3$^+$ Tregs **A)** and Flu-sp CD8+ T cells **B)** in spleens of infected mice. **C), D)** Representative flow cytometric analysis and quantification of expression of the indicated markers by lung parenchymal **D)** or blood exposed Flu-sp CD8 T cells. **E)** Total NP-specific IgG antibody measured by ELISA at the indicated serial dilutions of serum collected at necropsy in naïve and vehicle- or Fc.Mut24-treated mice as indicated. **F)** Representative flow cytometry analysis of CD4 MHCII tetramer against $NP_{311\text{-}325}$ staining and quantitative analysis of flu-sp CD4$^+$ T cells in the lung (left) and spleen (right) of vehicle- and Fc.Mut24-treated mice. **G)** Quantification of FoxP3 negative CD4 T cells in lung and spleen. **H)** Quantification of IFN-γ production after stimulation. **I)** Quantification of IL-2 production after stimulation. Representative of two separate experiments n = 10, except F which n = 5. $*p < 0.05$, $**p < 0.01$, $****p < 0.0001$, by unpaired T test for all panels except E where a one way ANOVA with multiple comparison was used.

### Supplemental Data Figure 2

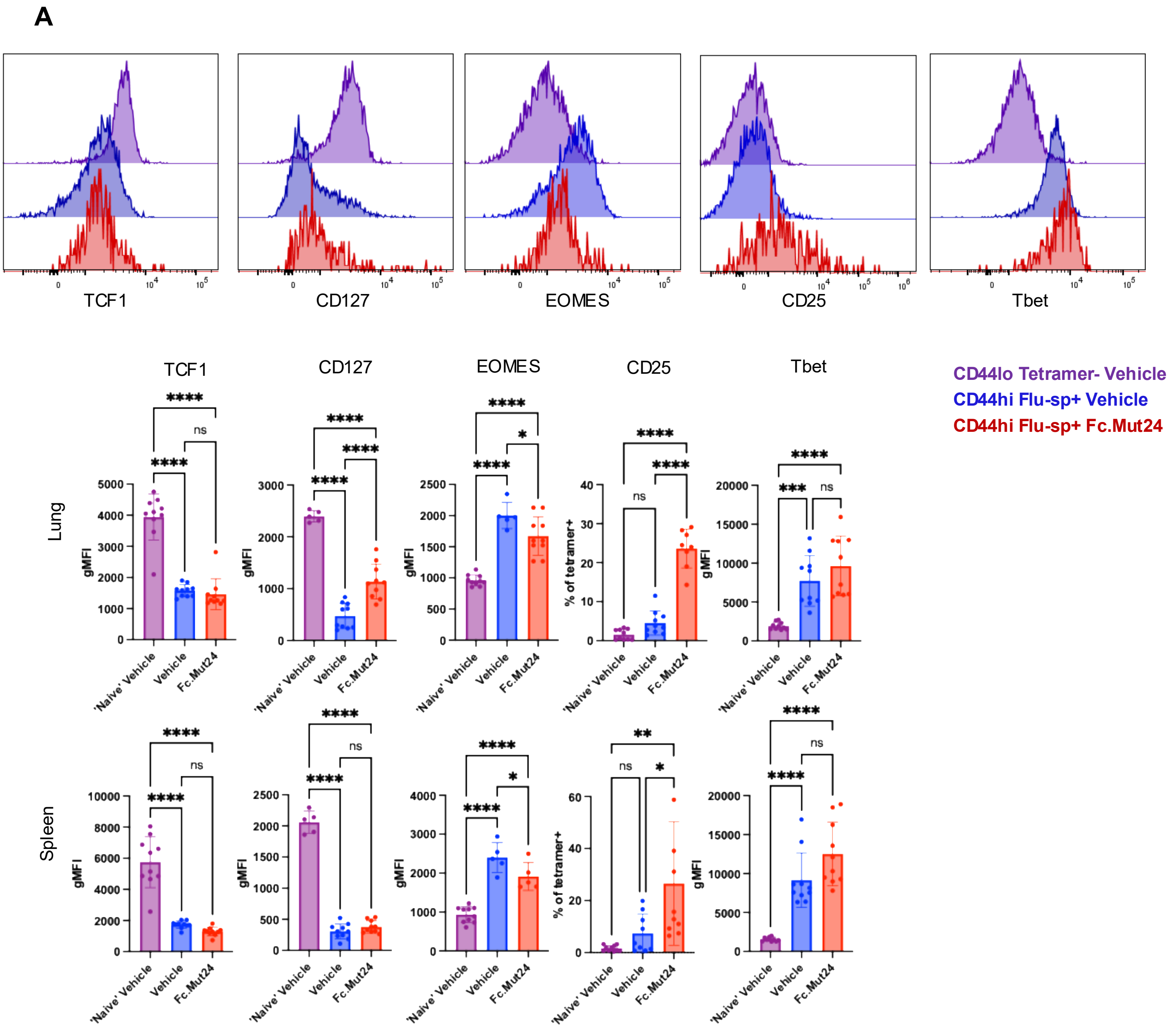


**Supplemental Figure 2**. Mice were with vehicle or Fc.Mut24 one day before infection with 2000PFU of X31 **A)** Representative marker staining and quantitative analyses of expression of the indicated makers by gated CD44loTetramer- naïve T cells from vehicle-treated mice, and from Flu-sp T cells from vehicle- (blue) or Fc.Mut24-treated (red) mice in the lungs (top) and spleen (bottom). Representative of two separate experiments n = 10 *p < 0.05, **p < 0.01, ****p < 0.0001, by one way ANOVA with multiple comparison.

### Supplemental Figure 4

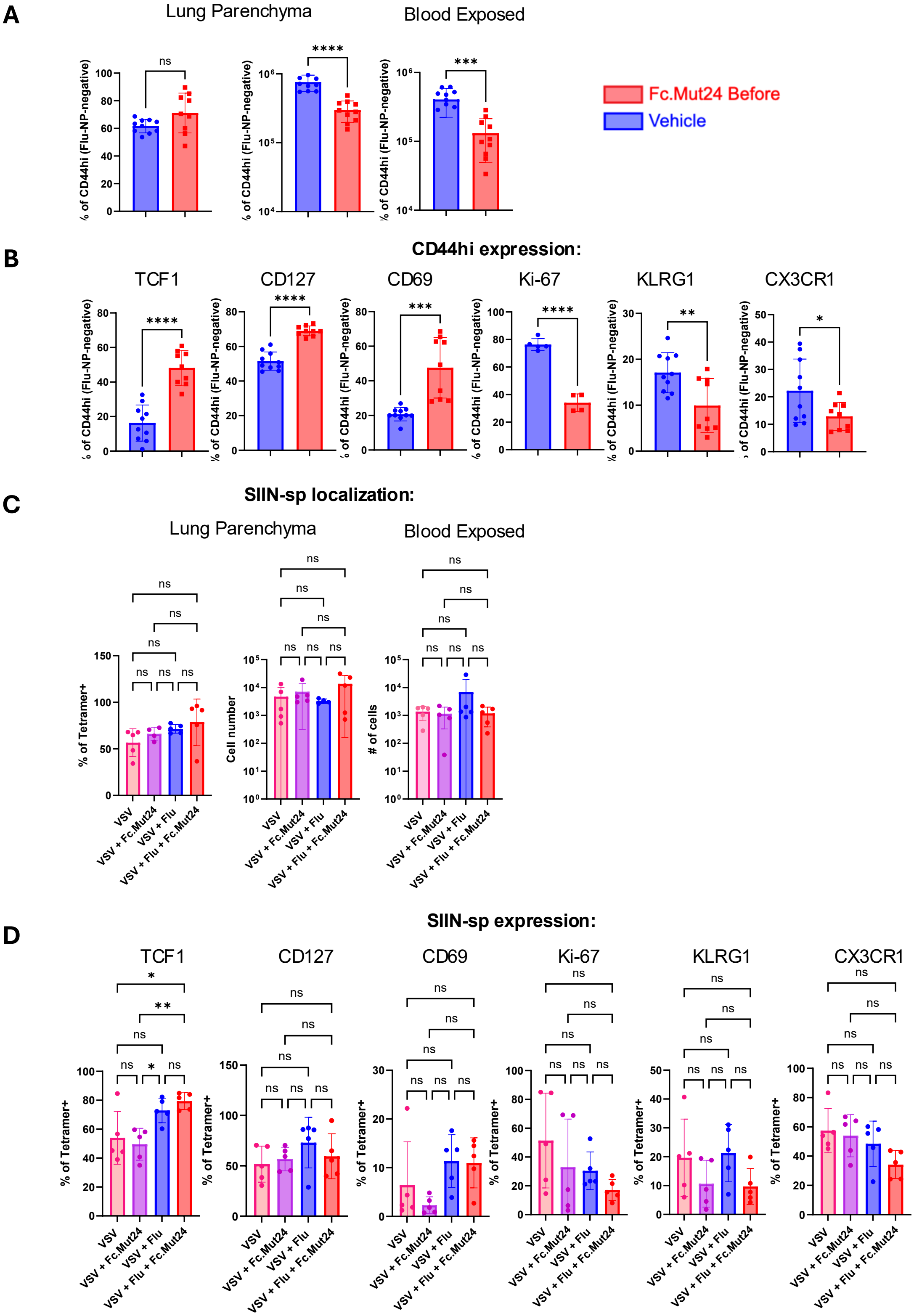


**Supplemental Figure 3. A)** Quantification of CD44hi Tetramer- CD8 T cells based on IV fractions in the lungs of mice. **B)** Quantification of selected markers on CD44hi Tetramer- CD8+ T cells. **C)** Quantification of bystander SIIN-sp CD8 T cells in lung based on IV labeling **D)** Quantification of indicated markers on SIIN-sp CD8 bystanders. Panels A and B represent two separate experiments n = 10. C and D are representative of one experiment n = 5 *p < 0.05, **p < 0.01, ****p < 0.0001, by unpaired T test for panels A and B and by one way ANOVA with multiple comparison for panel C and D.

### Supplemental data Figure 5

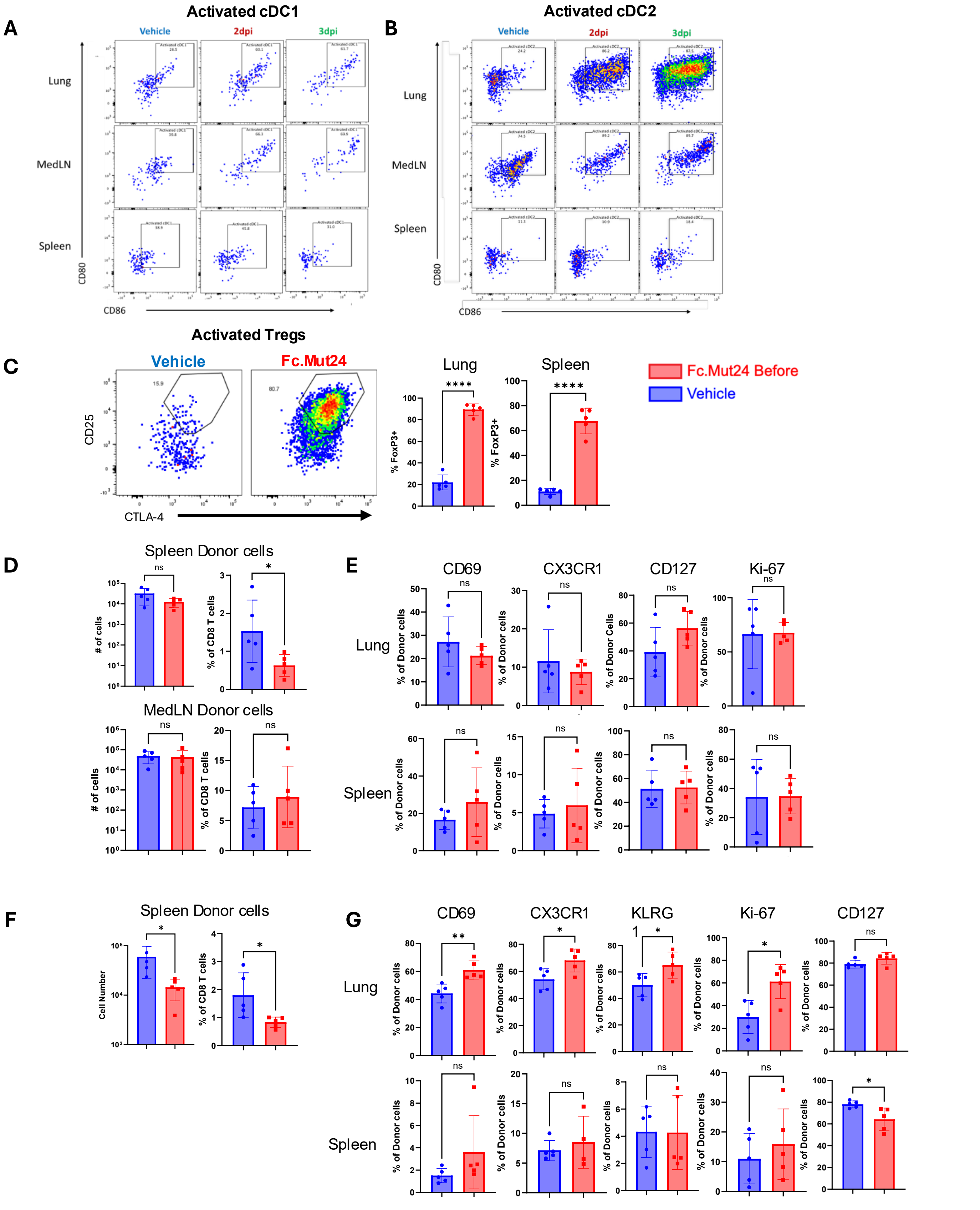


**Supplemental Figure 4**. Mice were treated with vehicle (blue) or Fc.Mut24 (red) one day before infection with 2000PFU of X31. **A), B)** An analysis of CD80 and CD86 expression by cDC1 **A)** and cDC2 **B)** at 0, 2 or 3dpi as indicated. **C)** Representative flow cytometry and quantitative analysis of CD25 and CTLA4 expression by gated CD4+Focp3+ Treg in vehicle- or Fc.Mut24-treated mice at 3dpi (right). **D), E)** Analysis of donor OT-1 T cells given to mice before X31-SIINFEKL infection. Quantification of transferred OT-1 cells in the spleen and medLN of vehicle- and Fc.Mut24-treated mice at 4dpi D), expression of indicated activation markers by donor OT-1 cells isolated from lungs (top) and spleen (bottom) at 4dpi. **E). F), G)** Analysis of donor OT-1 T cells given to mice at 3dpi with X31-SIINFEKL infection. Quantification of donor OT-1 cells that were transferred at 3dpi and from the spleen at takedown (9dpi) **F)**, expression of indicated activation markers by donor OT-1 cells isolated from lungs (top) and spleen (bottom) at 9dpi. Experiments representative of an experiment n = 5. $*p < 0.05$, $**p < 0.01$, $****p < 0.0001$, by unpaired T test for all panels.

### Supplemental Data 5

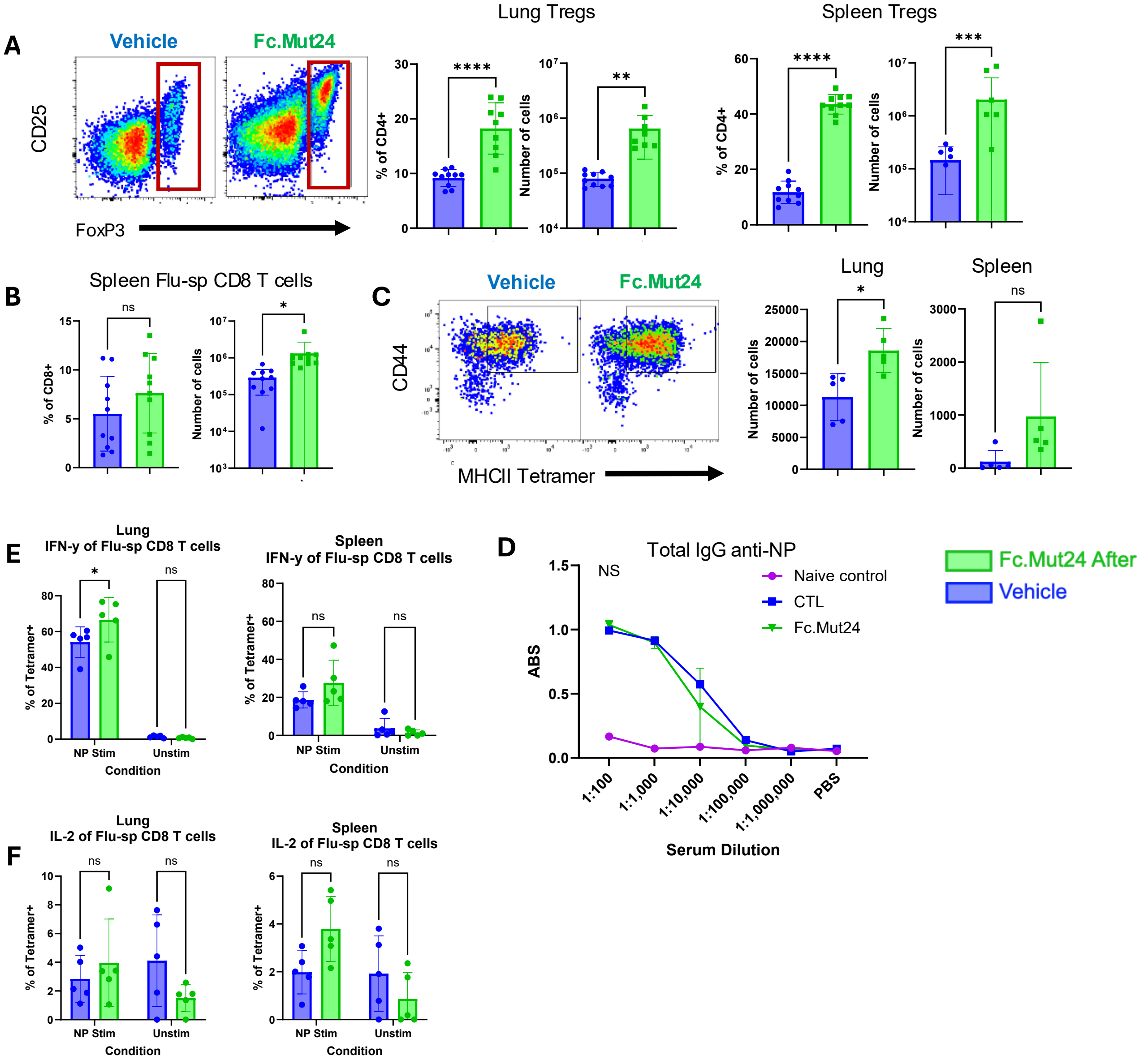


**Supplemental Figure 5.** Mice were treated with vehicle (blue) or Fc.Mut24 (green) at 4dpi with X31. All data represent takedown at 9dpi. **A)** Representative flow cytometry analysis of CD25 and FoxP3 expression by gated CD4+ T cells in the lungs of vehicle and Fc.Mut24 treated mice, and quantitative analysis of the frequency and number of Treg cells in the lung and spleen. **B)** Number and frequencies of Flu-sp CD8 T cells spleen of mice treated as indicated. **C) F)** Representative flow cytometry analysis of CD4 MHCII tetramer staining against NP311-325 and quantitative analysis of flu-sp CD4+ T cells in the lung (left) and spleen (right) of vehicle- and Fc.Mut24-treated mice. **D)** Total NP-specific IgG antibody measured by ELISA at the indicated serial dilutions of serum collected at necropsy in naïve and vehicle- or Fc.Mut24-treated mice as indicated. **E)** Quantification of IFN-γ production after stimulation. **F)** Quantification of IL-2 production after stimulation. All data represent two separate experiments n = 10, except for C where n = 5. *$p < 0.05$, **$p < 0.01$, ****$p < 0.0001$, unpaired T test for all panels except for D where a one way ANOVA with multiple comparison was used.

### Supplemental data 6

**A** Expression of markers on Flu-sp CD8 T cells

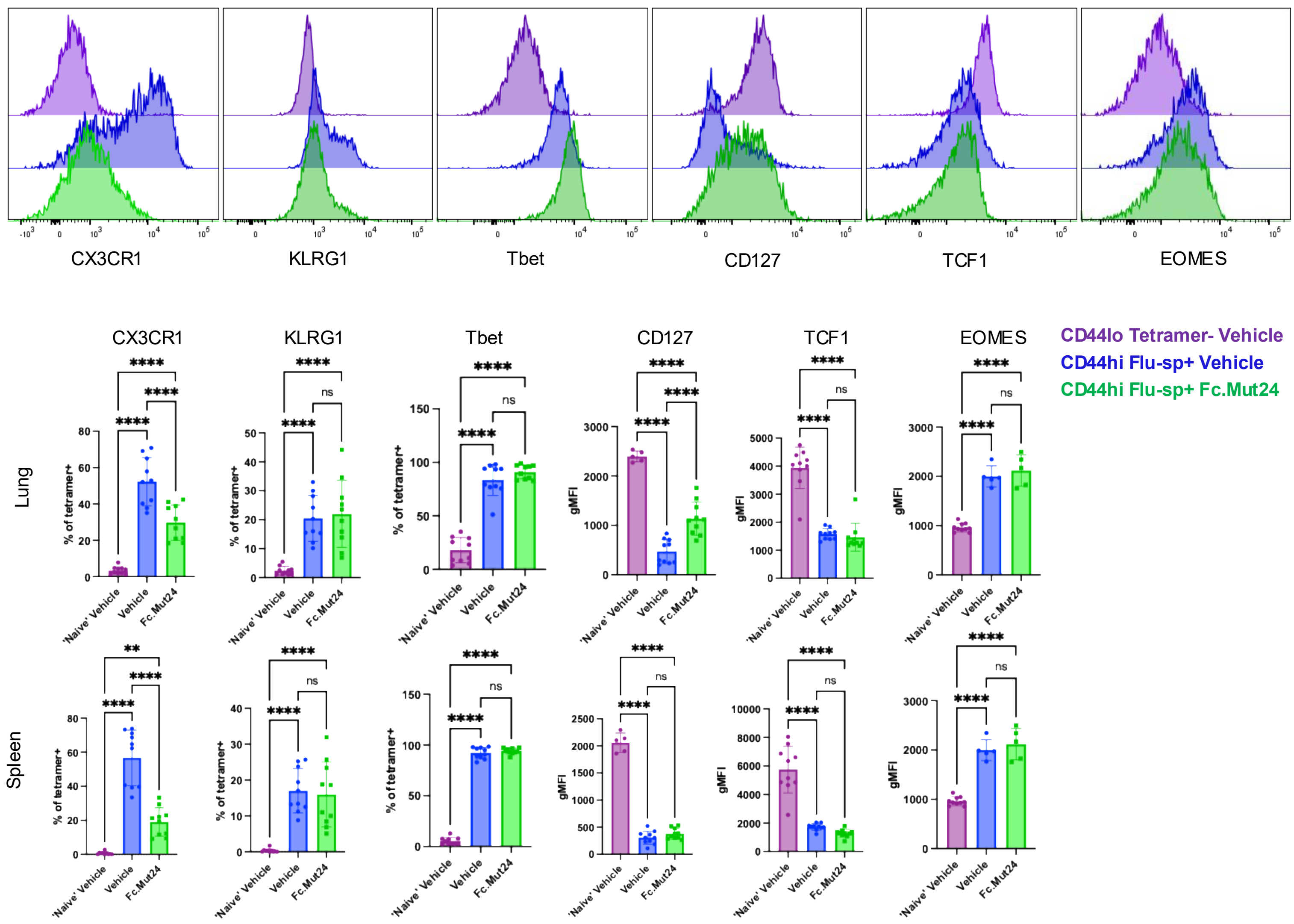


**Supplemental Figure 6**. Mice were treated with vehicle or Fc.Mut24 at 4dpi with X31. All data represent takedown at 9dpi. A) Representative flow cytometry analysis of the indicated marker expression by gated CD44loTetramet- naïve T cells from vehicle-treated mice (purple), or by Flu-sp CD8+ T cells from lungs and spleens of vehicle- (blue) or Fc.Mut24-treated (green) mice, and quantification of marker expression by cells from the lungs and spleen as indicated. Representative of two separate experiments n = 10 *p < 0.05, **p < 0.01, ****p < 0.0001, by one way ANOVA with multiple comparison.

### Supplemental Figure 7

**A** CD25 Frequencies on Total CD8+ T cells post 48hr activation

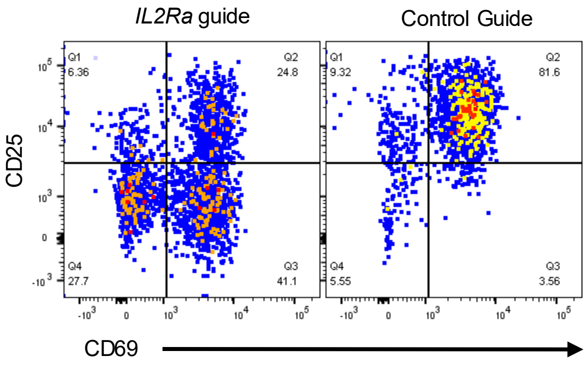


**Supplemental Figure 7. A**) Representative flow cytometry analysis of CD25 and CD69 expression by OT-1 T cells following transduction of CAS9 RNPs containing control or Il2ra-targeting guide RNA following 48h activation with plate bound anti-CD3/CD28.

### Supplemental Figure 8

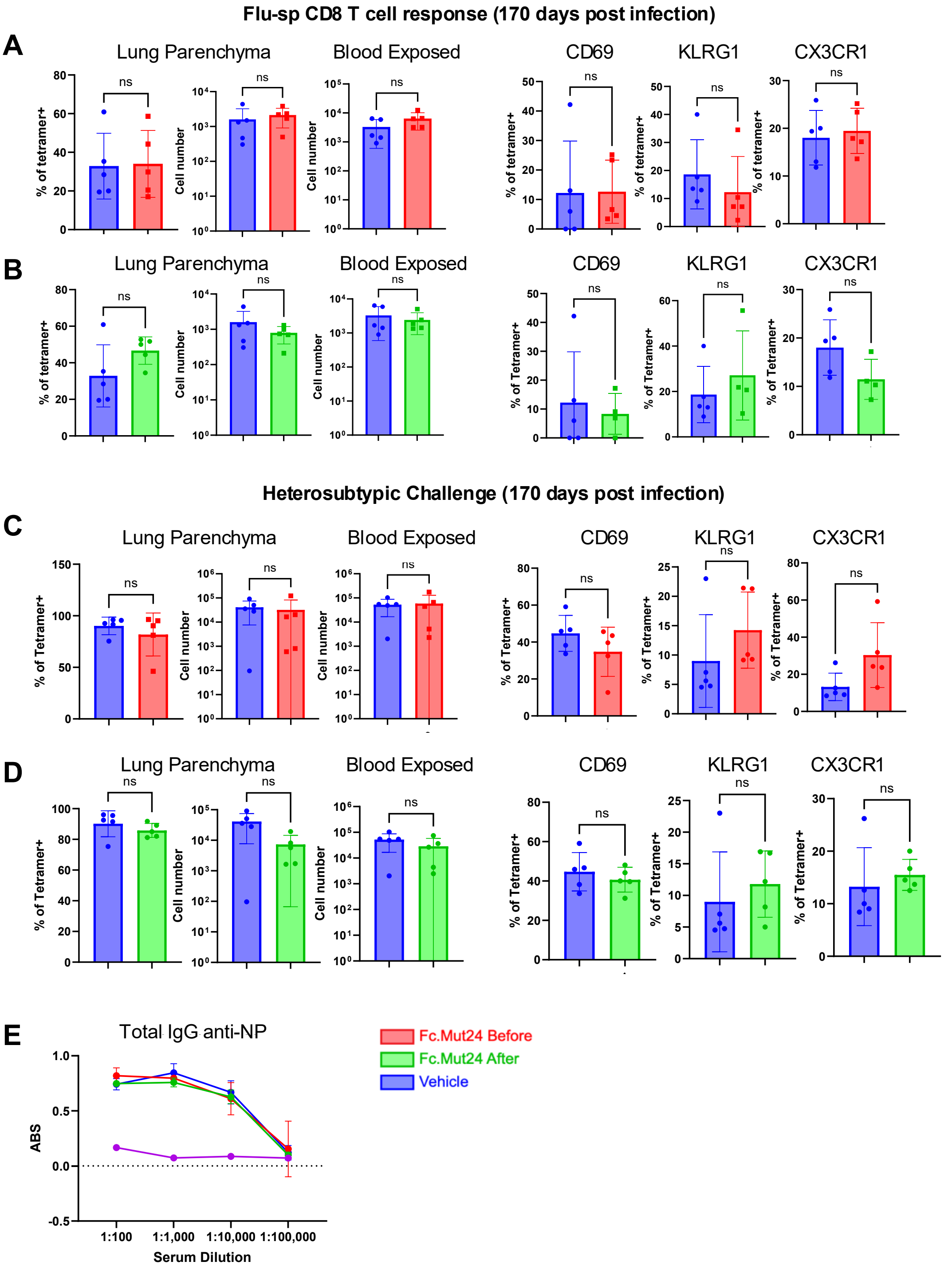


**Supplemental Figure 8.** Mice were all treated a day prior to (**A and B**) or 4 days after infection with X31 with Fc.Mut24 (**C and D**). **A or C)** Markers on Flu-sp CD8 t cells in the lung of mice treated with Fc.Mut24 before or after (respectively) X31 at 170dpi. **B or D)** Markers on Flu-sp CD8 t cells in the lung of mice treated with Fc.Mut24 before or after (respectively) X31 at 170dpi and then lethally challenged with 5000 PFU of PR8. **E)** total IgG titers against the Flu NP from mice prior to 170dpi. Representative of one experiment where n = 5 *$p < 0.05$, **$p < 0.01$, ****$p < 0.0001$, by unpaired T test.
